# Measuring conformational equilibria in allosteric proteins with time-resolved tmFRET

**DOI:** 10.1101/2023.10.09.561594

**Authors:** William N. Zagotta, Eric G. B. Evans, Pierce Eggan, Maxx H. Tessmer, Kyle D. Shaffer, E. James Petersson, Stefan Stoll, Sharona E. Gordon

## Abstract

Proteins are the workhorses of biology, orchestrating a myriad of cellular functions through intricate conformational changes. Protein allostery, the phenomenon where binding of ligands or environmental changes induce conformational rearrangements in the protein, is fundamental to these processes. We have previously shown that transition metal Förster resonance energy transfer (tmFRET) can be used to interrogate the conformational rearrangements associated with protein allostery and have recently introduced novel FRET acceptors utilizing metal-bipyridyl derivatives to measure long (>20 Å) intramolecular distances in proteins. Here, we combine our tmFRET system with fluorescence lifetime measurements to measure the distances, conformational heterogeneity, and energetics of maltose binding protein (MBP), a model allosteric protein. Time-resolved tmFRET captures near-instantaneous snapshots of distance distributions, offering insights into protein dynamics. We show that time-resolved tmFRET can accurately determine distance distributions and conformational heterogeneity of proteins. Our results demonstrate the sensitivity of time-resolved tmFRET in detecting subtle conformational or energetic changes in protein conformations, which are crucial for understanding allostery. In addition, we extend the use of metal-bipyridyl compounds, showing Cu(phen)^2+^ can serve as a spin label for pulse dipolar electron paramagnetic resonance (EPR) spectroscopy, a method which also reveals distance distributions and conformational heterogeneity. The EPR studies both establish Cu(phen)^2+^ as a useful spin label for pulse dipolar EPR and validate our time-resolved tmFRET measurements. Our approach offers a versatile tool for deciphering conformational landscapes and understanding the regulatory mechanisms governing biological processes.

**STATEMENT OF SIGNIFICANCE:** Responsible for the regulation of virtually all biological processes, allosteric proteins are fundamental to all life. To understand the mechanisms for their vital functions, we must determine the structure and energetics of these proteins under physiological conditions. In this work, we have developed a fluorescence method that allows for the simultaneous measurement of protein structure and energetics under physiological conditions. These studies pave the way for future advances in physiology and medicine.

## INTRODUCTION

Protein allostery plays a pivotal role in the regulation of virtually all biological processes. In response to the binding of specific molecules or changes in environmental conditions, allosteric proteins undergo distinct changes in structure that regulate the protein’s activity or interaction with other molecules. This dynamic behavior allows proteins to function as molecular switches, orchestrating a wide range of biological functions such as enzymatic catalysis, signal transduction, gene regulation, and cellular motility.

The mechanism of allostery involves a choreography of the protein’s structure and energetics. A stereotypical allosteric protein might have two conformations, a resting state and an active state (Figure 1A). Whereas the active state might be energetically unfavorable (positive Δ*G*) in the absence of ligand, it becomes more favorable (negative Δ*G*) in the presence of ligand. In this way, the conformation of the protein, and therefore its activity, is coupled to the binding of a ligand.

**Figure 1.**
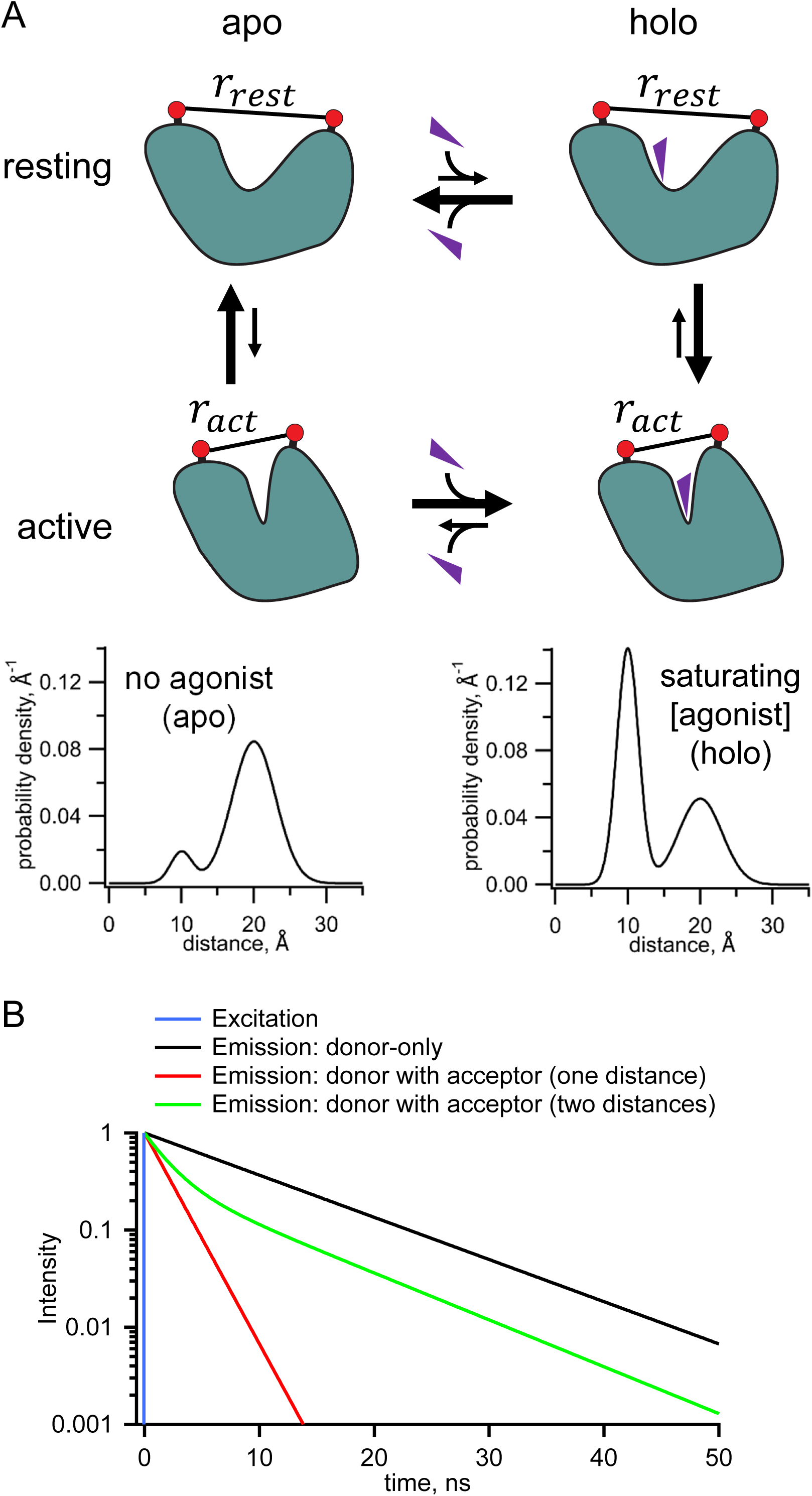
Measurement of distance distributions for an allosteric protein. (**A**) Conformational states (top) and distance distributions (bottom) for a stereotypical allosteric protein. The red circles represent the FRET probes and the purple triangles the ligand. (**B**) Theoretical effect of FRET with one or two distances on the fluorescent lifetimes in the time domain.

One general way to determine this Δ*G* is to measure conformational equilibria based on distance distributions between a pair of probes, attached to the protein of interest, which undergo a change in separation between resting and active states (Figure 1A). The hypothetical distance distributions in Figure 1A show that, in the absence of ligand, or “apo” condition, the resting conformation (longer distance) dominates, and, in saturating ligand or “holo” condition, the active conformation (shorter distance) dominates. The relative occupancy of the two conformations can be quantified by the relative area of the probability distribution for each conformational state. The ratio of the areas reveals the equilibrium constant and therefore the ΔG for the transition between the apo and holo states. These distance distributions reveal two types of heterogeneity that must be considered for any allosteric protein: 1) heterogeneity in functional state, as both resting and active conformations are present in both the absence and presence of ligand, and 2) for any given state, heterogeneity in the protein backbone and probe rotameric ensembles produce a distribution of distances between probes.

While Förster resonance energy transfer (FRET) has been used as a “molecular ruler” to measure distances in proteins, standard steady-state FRET experiments provide just a single number, the apparent FRET efficiency, from which one can calculate only a single weighted-average distance (1,2). Time-resolved FRET experiments, however, generate richer data, from which the distribution of distances can be recovered (2–9). Time-resolved FRET utilizes fluorescence lifetimes, the latencies between absorption and emission of photons from a fluorophore. In the simplest cases, fluorescence lifetimes are single exponentially distributed with a time constant of a few nanoseconds. The presence of a FRET acceptor accelerates the fluorescence decay of the donor fluorophore in a manner that is highly dependent on the distance between the donor and acceptor (Figure 1B). If, for example, there are two states with different distances between the donor and acceptor, the decay will be double exponential with the fraction of each component representing the prevalence of that state. Importantly the interconversion among states is generally slower than the nanosecond fluorescence lifetime, so time-resolved FRET captures a near-instantaneous snapshot of the distance distribution without the averaging of distances, in contrast to steady-state or single-molecule fluorescence methods.

The utility of time-resolved FRET for deciphering protein dynamics has been limited by several experimental factors: 1) Labeling proteins site-specifically with donor and acceptor fluorophores can be a challenge. 2) The large size of most visible-light fluorophores and the length of the linkers that connect the fluorophores to the protein make it difficult to discriminate backbone dynamics and energetics from those of the fluorophore/linker. 3) Most fluorophores demonstrate multi-exponential lifetimes, which makes analysis of FRET data more complicated. 4) The fluorescence lifetimes of most dyes in common use are in the few-nanosecond range, giving a limited dynamic range for measuring time-resolved FRET. We recently developed a novel system for time-resolved FRET that overcomes these limitations by combining a noncanonical amino acid fluorophore donor and a transition metal ion acceptor (10). Here we employ this novel system with metal-bipyridyl acceptors, developed in a companion paper in this issue (11), to measure longer distance distributions in a model protein, maltose binding protein (MBP). We show that time-resolved FRET can quantify both the heterogeneity of a given conformational state and the energetics that govern the distribution of a protein among conformational states, collectively referred to as protein dynamics.

## MATERIALS AND METHODS

### Expression and purification of MBP

For tmFRET experiments, the expression and purification of MBP was done as described previously (11). Briefly, MBP TAG constructs with a C-terminal twin-strep tag were cotransfected with a plasmid containing the AcdA9 aminoacyl tRNA synthetase and its cognate tRNA (12) in BL-21(DE3) cells. Cultures were induced in the presence of 0.6 mM Acd in the media, and MBP was purified on a Streptactin column (IBA Life Sciences, Göttingen, Germany) column.

For RIDME experiments, dual cysteine constructs of MBP-295C-211C and MBP-322C-278C with N-terminal 6×His tags were expressed from a pETM11 vector in *E. coli* C43(DE3) and subsequently purified by Co^2+^ affinity chromatography as previously described (13). The 6xHis tag was removed by incubation with a 1:50 (TEV:MBP) weight ratio of TEV protease (4 h at room temperature, then 12 h at 4 °C) in K^+^-Tris buffer (130 mM KCl, 30 mM Tris, pH 7.4) containing 0.5 mM EDTA and 1 mM TCEP. The reaction was desalted into K^+^-Tris buffer (pH 7.4) plus 5 mM imidazole and 50 μM TCEP and further purified by reverse IMAC over TALON resin. Flow through containing cleaved MBP was supplemented with 1 mM TCEP and 5 mM EDTA, concentrated (30 kDa MWCO), and stored at 4 °C.

### Labeling reagents

PhenM and [Ru(bpy)_2_phenM]^2+^ were prepared as stocks in DMSO and used within minutes of final dilution into aqueous solution. 2 M hydroxylamine hydrochloride in water was prepared for use in Fe^2+^ experiments and used for only one day. FeCl_2_ was prepared as a 100 mM stock with 1 M hydroxylamine hydrochloride in water and made fresh for each experiment day. CuCl_2_ was prepared as a 100 mM stock in water.

### [Ru(bpy)2phenM]^2+^ labeling

Time-resolved fluorescence measurements required a higher protein concentration (i.e., higher concentration of donor) and therefore a higher concentration of [Ru(bpy)_2_phenM]^2+^ was required for labeling. To reduce background absorption due to [Ru(bpy)_2_phenM]^2+^ in solution, we first labelled with [Ru(bpy)_2_phenM]^2+^ and then column purified the protein. Specifically, 100 mM [Ru(bpy)_2_phenM]^2+^ stock in DMSO was diluted to 1 mM in the concentrated protein in K^+^-Tris buffer (pH 7.4). After 10 minutes, the solution was passed over a Bio-Rad Micro Bio-Spin 6 column that had been equilibrated with K^+^-Tris buffer (pH 7.4) to remove unreacted label. K^+^-Tris buffer (pH 7.4) was also used to elute the protein.

### [Fe(phenM)_3_]^2+^ labeling

For [Fe(phenM)_3_]^2+^ experiments, a solution of 920 µM FeCl_2_ in 9.2 mM hydroxylamine hydrochloride with 2.3 mM phenM was prepared in water. After recording the donor-only fluorescence lifetime, this [Fe(phenM)_3_]^2+^ solution was added to the protein drop to a final concentration of 76.8 µM Fe^2+^, 768 µM hydroxylamine hydrochloride, and 192 µM phenM. For these brief experiments, additional hydroxylamine hydrochloride was not required to prevent oxidation of Fe^2+^ to Fe^3+^.

### [Fe(phenM)]^2+^ labeling

For [Fe(phenM)]^2+^ experiments, a 20 mM phenM stock in K^+^-Tris buffer (130 mM KCl, 30 mM Tris, pH 8.3) was added to MBP protein to achieve a final concentration of 2 mM phenM. After 10 minutes, the solution was passed over a Bio-Rad Micro Bio-Spin 6 column that had been equilibrated with K^+^-Tris buffer (pH 8.3) to remove unreacted phenM label. K^+^-Tris buffer (pH 8.3) was also used to elute the protein. After measuring the lifetime of the purified protein in the absence Fe^2+^, we added Fe^2+^ (with a 10-fold excess of hydroxylamine hydrochloride) to a final concentration of 800 μM.

### Measurement of fluorescence lifetime using FLIM

The theory underlying our FRET measurements with fluorescence lifetimes is well described elsewhere (2). Briefly, FRET decreases the fluorescence lifetime of a donor fluorophore by providing an additional path by which an excited state electron can lose its energy. When using a pulse excitation source and measuring fluorescence in the time domain, the decrease in lifetime is readily apparent as a faster decay in fluorescence intensity after excitation (Figure 1B). When using a frequency (ω) modulated excitation source, the lifetimes of donor in the absence and presence of acceptor are determined from the phase delays (the phase shift between the excitation and emission, φ_*ω*_) and modulation ratios (the fractional decrease in the amplitude of the emission, *m*_*ω*_) at each frequency (c.f. Figure 2A). With our frequency domain instrument, the frequency dependence of both φ_*ω*_and *m*_*ω*_are required to resolve complex lifetimes. A similar analysis can be performed using a time domain instrument (2).

**Figure 2.**
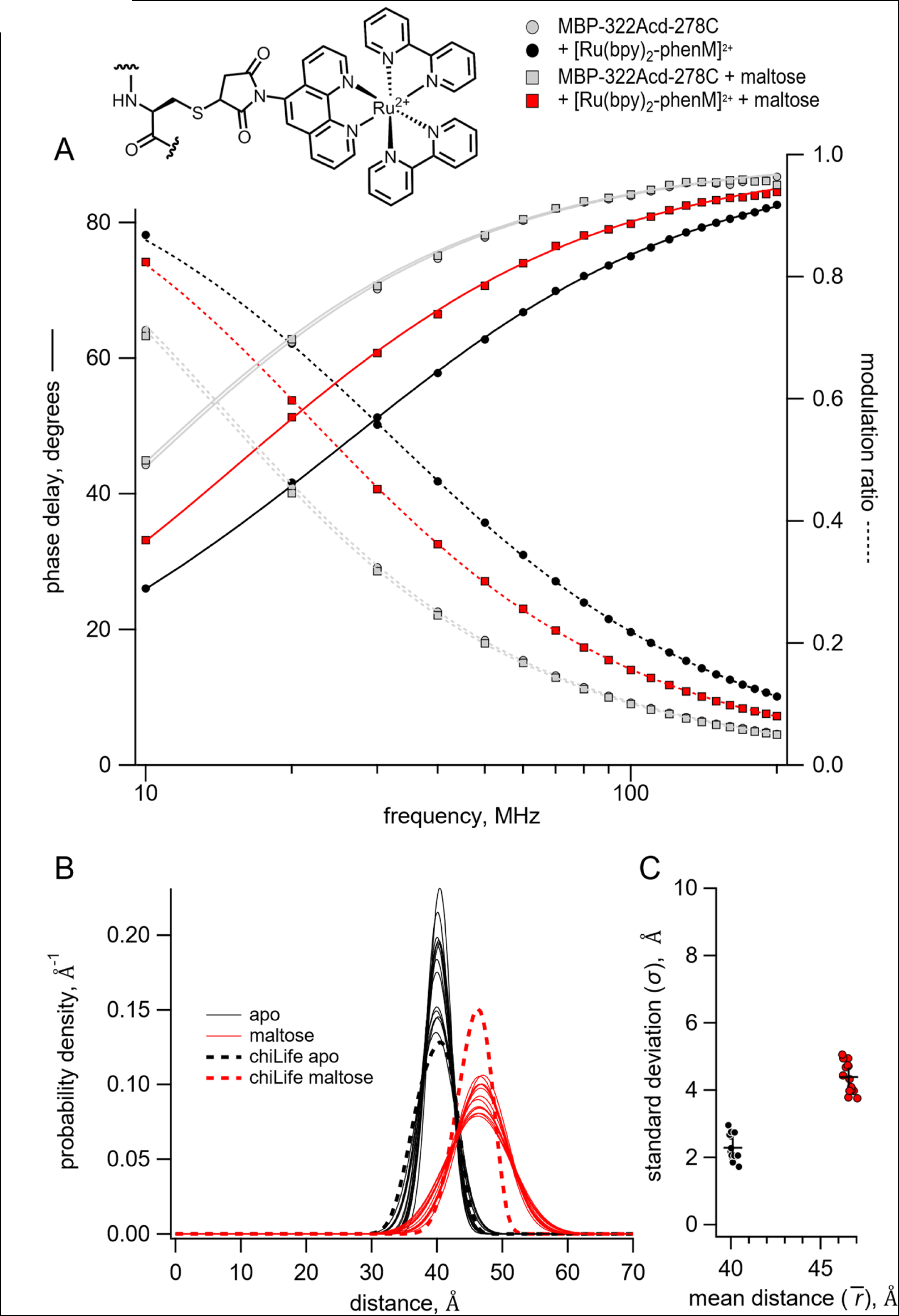
Time-resolved tmFRET of MBP-322Acd-278C labelled with [Ru(bpy)_2_phenM]^2+^. Structure of [Ru(bpy)_2_phenM]^2+^ label on the protein and legend for all panels are shown at the top. (**A**) Weber plot of the fluorescence lifetimes with 0 and 3 mM maltose in the presence and absence of [Ru(bpy)_2_phenM]^2^. (**B**) Spaghetti plot of the distance distributions measured by time-resolved tmFRET (thin solid curves) with 0 and 3 mM maltose (n=14) compared to the predicted distributions from chiLife for the resting and active state (dashed curves). (**C**) Scatter plot of the standard deviation vs. the mean distance for the Gaussian distributions with 0 and 3 mM maltose (n=14). The average values are shown as black +.

Frequency domain fluorescence lifetime data were collected using a Q2 laser scanner and A320 FastFLIM system (ISS, Inc., Champaign, IL, USA) mounted on a Nikon TE2000U microscope (Melville, NY, USA) and VistaVision software (ISS, Inc.). Acd or Atto 425 (the standard for calibration of the fluorescence lifetime) were excited using a 375 nm pulse diode laser (ISS, Inc.), driven by FastFLIM at the repetition rate of 10 MHz, with a 387 nm long-pass dichroic mirror, and emission was collected using a 451/106 nm band-pass emission filter and Hamamatsu model H7422P PMT detector. Affinity purified protein was used after about a 1:10 dilution in K^+^-Tris buffer. For each experiment, 11 μl of fluorescent sample was pipetted onto an ethanol-cleaned #1.5 glass coverslip mounted directly above the 10x 0.5 NA objective. Other reagents (maltose, [Ru(bpy)_2_phenM]^2+^, [Fe(phenM)_3_]^2+^, Fe^2+^, or EDTA) were pipetted directly into the sample drop and mixed at the final concentrations indicated in the text. For each condition, 256x256 confocal images were collected with a pinhole of 200 µm and a pixel dwell time of 1 ms. The pixels were averaged together for analysis, except as described for the phasor plot.

The experimental phase delays (φ_*ω*_) and the modulation ratios (*m*_*ω*_) of the fluorescence signal in response to an oscillatory stimulus with frequency ω were obtained using VistaVision software from the sine and cosine Fourier transform of the phase histogram *H(p)*, subject to the instrument response function (IRF) calibrated with 2 µM Atto 425 in water with a lifetime of 3.6 ns (2,14,15).

### Analysis of frequency domain fluorescence lifetime data

The theoretical estimates for φ_*ω*_and *m*_*ω*_were calculated from a model for fluorescence lifetime and FRET that assumes a single-exponential donor fluorescence lifetime with one or two Gaussian-distributed distances between the donor and acceptor as previously described with some modification (2,9,10,16,17). The phase delays (φ_*ω*_) and modulation ratios (*m*_*ω*_) were calculated as a function of the modulation frequency (ω) using the following equations:

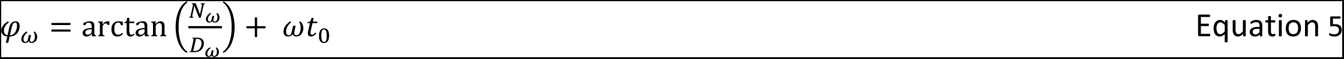

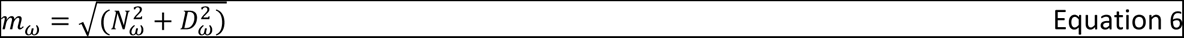

where the *N*_*ω*_corresponds to the out-of-phase component and *D*_*ω*_ corresponds to the in-phase component of fluorescence and *t*_0_. is the time shift of the IRF (Figure S1E). The two components for the fluorescence response with a contaminating background fluorescence were calculated using the following equations:

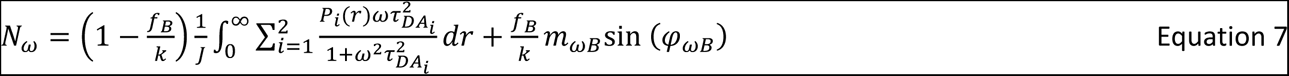

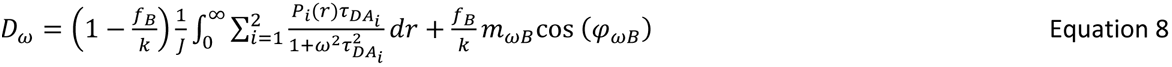

where *f*_B_is the fraction of the fluorescence intensity due to background (Figure S1D), and φ_ωB_ and *m*_ωB_ are the phase delay and modulation ratio of the background fluorescence measured from samples of K^+^-Tris Buffer without or with maltose. The normalization factor *J* is given by:

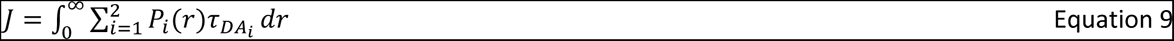

The decay time constant of the *i*th component of the donor lifetime in the presence of acceptor (τ_DAi_) is given by:

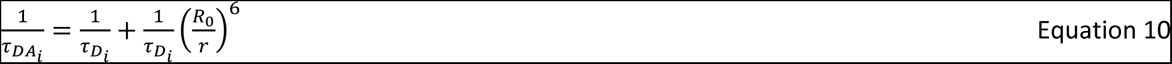

where τ_Di_ is the decay time constant of the donor in the absence of acceptor (Figure S1A), *r* is the distance between the donor and acceptor, and *R*_0_ is the characteristic distance for the donor-acceptor pair (Figure S1B).

The apparent FRET efficiency based on the donor quenching by the acceptor (*E*) was calculated using the following equation:

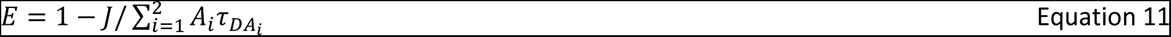

The correction factor for the fraction of background for the decrease in intensity due to FRET (*k*) was given by the following equation:

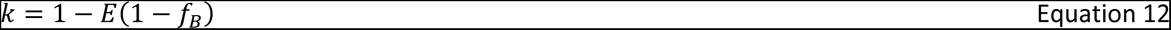

The distribution of donor-acceptor distances (*P*(*r*)) was assumed to be the sum of up to two Gaussians:

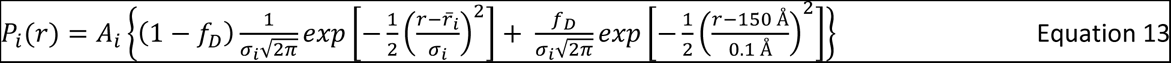

where *A*_i_, *r̅*_i_, and σ_i_ are the amplitude, mean, and standard deviation of the *i*th Gaussian respectively, and *A*_1_ + *A*_2_ = 1 (Figure S1C). The fraction donor only (*f*_D_) was modeled as a narrow Gaussian with a mean distance of 150 Å and a standard deviation of 0.1 Å, too far to exhibit any detectable FRET.

The phase delay and modulation ratios displayed were corrected for the background fluorescence and time shift in the IRF using the following equations:

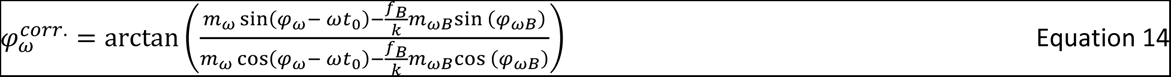

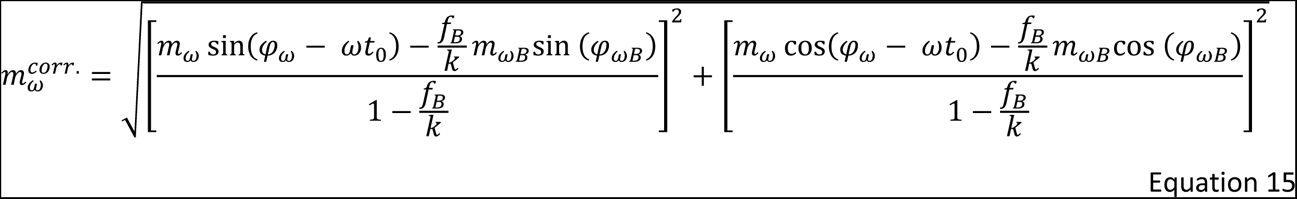

This Gaussian model for fluorescence lifetimes and FRET was implemented in Igor Pro v8 (Wavemetrics, Lake Oswego, OR) (code available at https://github.com/zagotta/FDlifetime_program_v2). The Gaussian model was globally fit, with *χ*^2^minimization, to the phase delay and modulation ratio for donor+acceptor fluorescence in the absence of maltose and in the presence of various concentrations of maltose as indicated in the text. There are 11 parameters in the parameter vector (*f*_D_, τ_D1_, τ_D2_, *R*_0_, *r̅*_1_, σ_1_, *A*_2_, *r̅*_2_, σ_2_, *t*_0_, *f*_B_) (Figure S1, blue variables). For the global fits. τ_D1_, τ_D2_, and *R*_0_ were constrained to previously determined values, *f*_D_, *r̅*_1_, σ_1_, *r̅*_2_, σ_2_, *f*_B_ were varied but constrained to be the same across all maltose concentrations, and *A*_2_ and *t*_0_ were varied separately for each maltose concentration. Occasionally, if *f*_D_,was inconsistent between individual fits of the same experiment, it was held at 0.1 as previously determined (11). Under these conditions, the values of the parameters were well determined (Figure S2) and independent of the initial guesses (Figure S3). Our donor fluorophore was determined to have a single exponential decay lifetime in our MBP constructs. However, the lifetime varied slightly in the absence and presence of maltose. The apo donor-only data were fit with *f*_D_ = 1 and *A*_2_ = 0 and three free parameters (τ_D1_, *t*_0_, *f*_B_), and the holo donor-only data were fit with *f*_D_ = 1 and *A*_2_ = 1 and three free parameters (τ_D2_, *t*_0_, *f*_B_) with. The τ_Di_ values for the [Ru(bpy)_2_phenM]^2+^ and [Fe(phenM)_3_]^2+^ experiments were averaged for 6-20 different samples and the average values used in the analysis for all experiments.

### In silico labeling and distance distribution simulations

Computational modeling of Acd and metal-phenM labels, as well as distance distribution predictions, were performed using chiLife (18) with the accessible-volume sampling method (19,20). Acd, and cysteine conjugates of [Ru(bpy)_2_phenM]^2+^, [Fe(phenM)_3_]^2+^, [Fe(phenM)]^2+^, and [Cu(phenM)]^2+^ were added as custom labels in chiLife. Briefly, starting label structures were constructed in PyMOL and energy minimized with the GFN force field (GFN-FF) in xTB (21). Custom labels were superimposed onto labeling sites of the target pdb structure, and mobile dihedral angles were uniformly sampled. Rotamers with internal clashes (< 2 Å) were discarded. External clashes were evaluated using a modified, repulsive-only Lennard-Jones potential and used to weight rotamers as previously described (19). The lowest weighted rotamers cumulatively accounting for a fraction of 0.005 of the total rotamer weights were discarded. Sampling was terminated once 10,000 samples had been attempted, generating between 400 and 2,500 rotamers, depending on the specific label and protein site. To calculate a simulated distance distribution between two rotamer ensembles, a weighted histogram was made for pairwise distances between the spin or fluorescent centers of each pair of rotamers from the two ensembles. For Acd, the center coordinates were defined by the mean position of all atoms in the central acridone ring. For the metal-phenM labels, the center coordinates were on the transition metal ion. Histograms were then convolved with gaussian distributions with a 1 Å standard deviation, and the resulting distributions were normalized.

### [Cu(phenM)]^2+^ spin-labeling and EPR sample preparation

Purified MBP-295C-211C and MBP-322C-278C (∼ 50 μM) were desalted (G-25) into K^+^-Tris buffer (pH 7.4) and immediately reacted with 0.5 mM phenM, freshly prepared from DMSO stock as a 5 mM solution in K^+^-Tris buffer with 1 mM EDTA. The reaction was nutated at 4 °C for 1 h, desalted (G-25) into K^+^-Tris buffer (pH 7.4), and concentrated (5 kDa MWCO). phenM-labeled MBP solutions were then incubated with 1 mM CuSO_4_ for 10 minutes at room temperature and loaded into 10 kDa MWCO microdialysis units (Thermo) and dialyzed against K^+^-Tris buffer prepared in deuterium oxide (D_2_O). Dialysis was carried out for ∼ 18 h, replacing dialysis buffer with fresh deuterated K^+^-Tris buffer twice. RIDME samples were prepared with ∼ 10 μM labeled MBP supplemented with 30 % (v/v) d8-glycerol. Holo MBP samples were additionally supplemented with 5 mM maltose from a stock solution in D_2_O. Samples were loaded into 1.5 mm OD/1.1 mm ID quartz tubes (Sutter) with flame-sealed bottoms and flash frozen in liquid nitrogen (LN2). Samples were stored at −80 °C until measurement. Samples for CW EPR were prepared similarly, but without use of deuterated buffers and with 25 % (v/v) glycerol. CW EPR samples were loaded into 4 mm OD quartz EPR tubes (Wilmad), frozen in LN2, and measured on the same day.

### EPR measurements

Continuous-wave EPR spectra were recorded at 112 K on a Bruker EMX spectrometer operating at X-band frequency (∼9.3 GHz) with a Bruker ER 4102SHQE resonator. Spectra were recorded with 100 kHz field modulation with a sweep rate of 3.6 G/s and a modulation amplitude of 5 G. Spectra were background subtracted and baseline corrected in LabVIEW™. For visual comparison between samples (Figure S4), spectra were normalized by the double integral of the respected field-modulated spectrum. Magnetic parameters g and A were determined by least-squares fitting spectra using EasySpin 6.0 (22), assuming axial g and A tensors and including anisotropic line broadenings as additional fit parameters (Figure S4).

Pulse EPR experiments were performed at Q-band frequency (∼34 GHz) using a Bruker EleXsys E580 spectrometer with an overcoupled Bruker EN 5107D2 resonator. Pulses were amplified with a 300 W TWT amplifier (Applied Systems Engineering) and sample temperatures of 20 K or 10 K were maintained using a variable-temperature cryogen-free system (Bruker/ColdEdge). RIDME was performed using the established 5-pulse sequence (*π*/2) ― τ_1_ ― (*π*) ― τ_1_ + *t* ― (*π*/2) ― *T*_R_ ― (*π*/2) ― (τ_2_ – *t*) ― (*π*) ― τ_2_ ― (echo) (23). *π* /2 and *π* pulses were 12 and 24 ns, respectively, and were applied at frequency and magnetic field values corresponding to the maximum of the Cu^2+^ echo-detected field swept spectrum. To avoid dynamic decoupling artifacts in the RIDME time-trace, τ_1_ was chosen to be longer than τ_2_, with values of 4 μs and 3.5 μs for τ_1_ and τ_2_, respectively (24). Echo modulations due to solvent deuterium were suppressed by averaging over τ_1_ and τ_2_ with 16 ns increments over 8 steps. The relaxation interval *T*_R_ was 195 μs and was selected to be ∼ 0.75 that of the spin-lattice relaxation time (*T*_1e_) of the Cu^2+^ spin label at 20 K determined by inversion recovery experiments. Echo crossings were removed with a 32-step phase cycle (25); however, a small echo crossing artifact at *t* ≈ 0 could not be removed by phase cycling. This artifact, along with residual ESEEM contributions, were removed by recording a second RIDME time-trace with identical pulse lengths and delays at 10 K, where *T*_R_ ≈ 0.04*T*_1e_. Division of the 20 K data sets by the 10 K data sets gave the artifact-free dipolar evolution time traces used in all distance distribution analyses (Figure S5).

Divided and phase-corrected RIDME time-traces were analyzed with DeerLab version 1.1 (26) using Tikhonov regularization and compactness regularization (27). Residual intermolecular background was simultaneously modeled with the RIDME foreground using a homogenous 3-dimensional background model. The dipolar kernel was modified by replacing the default free electron g-value with an effective g-value of 2.1203 for each Cu^2+^ ion, which was determined from fits to the dual labeled MBP-[Cu(phenM)]^2+^ CW EPR spectra. Uncertainty estimations were determined by the asymptotic method in DeerLab and 95% confidence intervals were plotted as error bands on the RIDME probability distributions. All plots were generated in KaleidaGraph version 5.0 (Synergy Software) and visualized with KaleidaGraph and Inkscape version 1.2.

## RESULTS

Time-resolved FRET can be measured in either the time domain, usually with time correlated single-photon counting (TCSPC), or in the frequency domain, with both approaches yielding equivalent information (2). Here, we measured fluorescence lifetimes using a frequency-domain lifetime instrument (see Materials and Methods). For frequency-domain measurements, the lifetime data are visualized with a Weber plot that shows the phase shift of the response (phase delay) and the decrease in the amplitude of the response (modulation ratio) as a function of the modulation frequency of the excitation light (Figure 2A). These data can then be fit with models for the lifetimes that assume that they have single-exponential, multiexponential, or nonexponential decays.

To investigate the utility of time-resolved FRET to measure distance distributions, we incorporated the fluorescent noncanonical amino acid acridon-2-ylalanine (Acd) into MBP using amber codon suppression in bacteria as previously described (12,28,29). MBP is a clamshell-shaped protein that undergoes a significant closure of the clamshell upon binding maltose. For these experiments, we used two donor sites for specific incorporation of Acd, amino acid 295 at the outer lip of the clamshell and 322 on the back side of the clamshell. These donor fluorophore sites were then paired with single cysteine mutations for incorporation of transition metal chelates as FRET acceptors. Using MBP allowed us to test if time-resolved tmFRET could measure distances, distance distributions, and state energetics over a range of distances in a protein with a well-characterized structure and conformational rearrangement.

Acd incorporated at both donor sites (MBP-322Acd and MBP-295Acd) exhibited long, single-exponential fluorescence lifetimes similar to free Acd (Figure 2A and 5A, gray symbols). For Acd at 295 in wildtype MBP, the lifetime was nonexponential due to quenching by proximal Y307; therefore, all of our MBP-295Acd constructs also contain a Y307S mutation (referred to here as MBP-295Acd) (10,30). For both sites, the lifetimes were slightly longer in the presence of maltose (MBP-295Acd: apo, 14.7 ± 0.02 ns (n=13); holo 15.3 ± 0.1 ns (n=6); MBP-322Acd: apo, 15.4 ± 0.01 ns (n=20); holo 15.7 ± 0.01 ns (n=9)). This is likely due to a small change in the environment of the incorporated Acd in the presence of maltose and was factored into our subsequent analysis.

### Time-resolved FRET with [Ru(bpy)_2_phenM]^2+^ and [Fe(phenM)_3_]^2+^produce accurate average distances and narrow distance distributions

In the previous paper, we have shown that the cysteine-reactive metal chelate [Ru(2,2ʹ-bpy)_2_(1,10-phenanthroline-5-maleimide)]^2+^ ([Ru(bpy)_2_phenM]^2+^) can act as a long-distance tmFRET acceptor for Acd (11). [Ru(bpy)_2_phenM]^2+^ exhibits a substantial absorption in the visible range that overlaps with the emission spectrum of Acd, giving an *R*_0_, the distance producing 50% FRET efficiency, of 43.5 Å.

Consistent with its *R*_0_, labeling of MBP-322Acd-278C with [Ru(bpy)_2_phenM]^2+^ produced a substantial maltose-dependent decrease in steady-state Acd fluorescence indicating the presence of FRET between Acd and [Ru(bpy)_2_phenM]^2+^ (11). These steady-state FRET measurements, however, do not reveal the conformational heterogeneity in the sample.

To determine if time-resolved FRET could be used to measure distance distributions, we measured the fluorescence lifetimes of MBP-322Acd-278C labelled with [Ru(bpy)_2_phenM]^2+^. As shown in Figure 2A, labeling with [Ru(bpy)_2_phenM]^2+^ caused a substantial decrease in the average lifetime (manifesting as a shift in the phase delay and modulation ratio curves to *higher* frequencies in the Weber plot). Furthermore, subsequent addition of maltose increased the average lifetime (shifting the phase delay and modulation ratio curves to *lower* frequencies) reflecting an increase in the average distance between Acd at 322 and [Ru(bpy)_2_phenM]^2+^ at 278C, as predicted from structural modeling (see below). No change in lifetime was observed for MBP-322Acd without the cysteine mutation (data not shown; no native cysteines are present in MBP). These data suggest that time-resolved tmFRET could be used to reveal the intramolecular distance distributions in our samples.

To quantify the distance distributions from the fluorescence lifetime data, we fit the data with a model that predicts the lifetimes for a distribution of distances. The model assumes the following: 1) The fluorescence lifetimes of the donor-only protein (i.e., in the absence of acceptor) is single-exponentially distributed with a time constant τ_*D*_ (though τ_*D*_ can be different in the resting and active states). 2) The decrease in lifetime in the presence of the attached acceptor is due to a FRET mechanism with a known *R*_0_. 3) Donor and acceptor dipoles are randomly oriented relative to each other (*κ*^2^ = 2/3), a reasonable assumption when one member of the FRET pair is a metal ion (31). 4) There is only a single acceptor for each donor. 5) The distance distribution for each state can be approximated by a Gaussian distribution with a distinct mean distance and standard deviation. And 6) the distances do not change appreciably on the time scale of the fluorescence lifetime. Most of these assumptions can be experimentally verified in our sample. This model was globally fit to the phase delay and modulation ratio data across multiple conditions in the same experiment (apo, holo, and intermediate concentrations of ligand). The values of 10 to 12 free parameters for each experiment were determined using *χ*^2^ minimization. Because of how the distributions were parameterized, the fits generally had fewer free parameters than fitting with a sum of exponentials. This approach was pioneered in the 1970s, primarily by Steinberg and coworkers (32,33).

Global fits of the Gaussian model to the fluorescence lifetime data of MBP-322Acd-278C labelled with [Ru(bpy)_2_phenM]^2+^ in the apo and holo states are shown in Figure 2A. The values of the mean distances and standard deviations in the apo and holo states for 14 different experiments are shown in a spaghetti plot in Figure2B and a scatter plot in Figure 2C and were very reproducible. To compare our data to the predictions of structural modeling, rotameric ensembles of Acd and [Ru(bpy)_2_phenM]^2+^ were modelled onto crystal structures of apo and holo MBP (34,35) using the accessible-volume approach in chiLife (18), and used to predict distance distributions for MBP-322Acd-278C labeled with [Ru(bpy)_2_phenM]^2+^ (Figure 2B). In both the absence and presence of maltose, the distance distributions were remarkably similar to those predicted by chiLife. The average experimental distances and maltose-dependent delta distance were within 1 Å of those predicted by chiLife, and the widths of the distributions were also similar (though the experimentally determined width was consistently larger for the holo state). This similarity suggests that specific interactions between the probes and the protein (not accounted for by chiLife) do not play a significant role at these sites. Both the experimentally determined and predicted distance distributions between Acd and [Ru(bpy)_2_phenM]^2+^ were surprisingly narrow, indicating that [Ru(bpy)_2_phenM]^2+^ did not add appreciably to the measured heterogeneity under these conditions.

To determine the ability of tmFRET between Acd and [Ru(bpy)_2_phenM]^2+^ to resolve probability distributions that are a mixture of resting and active states, we performed lifetime experiments with subsaturating maltose concentrations applied to MBP-322Acd-278C labelled with [Ru(bpy)_2_phenM]^2+^, one just below the *K*_D_ and another just above the *K*_D_. The intermediate concentrations produced intermediate curves on the Weber plot (Figure 3A). To analyze the lifetimes in a model-independent way, we graphed the data on a phasor plot, a plot of the out of-phase versus the in-phase components of the fluorescence in frequency domain experiments (14). This plot revealed that the data for these intermediate concentrations fall on a line between the zero and saturating maltose concentrations, indicating that these intermediate maltose concentrations are a mixture of the same resting and active states produced by zero and saturating maltose (Figure 3B). We therefore performed global fits of the Gaussian model to the data on the Weber plot at all four maltose concentrations (0, 5 μM, 9.2 μM, and 3 mM), constraining the mean distance and standard deviation for each state to be the same for all conditions and allowing the fraction in the active state to vary (Fig 3C). The maltose dependence of the fraction of the active state nicely conformed to a binding isotherm, yielding an affinity of 6.3 μM, consistent with that previously measured using steady-state fluorescence (11) (Figure 3D). This is remarkable given that the difference in the average distance between the resting and active states is only about 6 Å.

**Figure 3.**
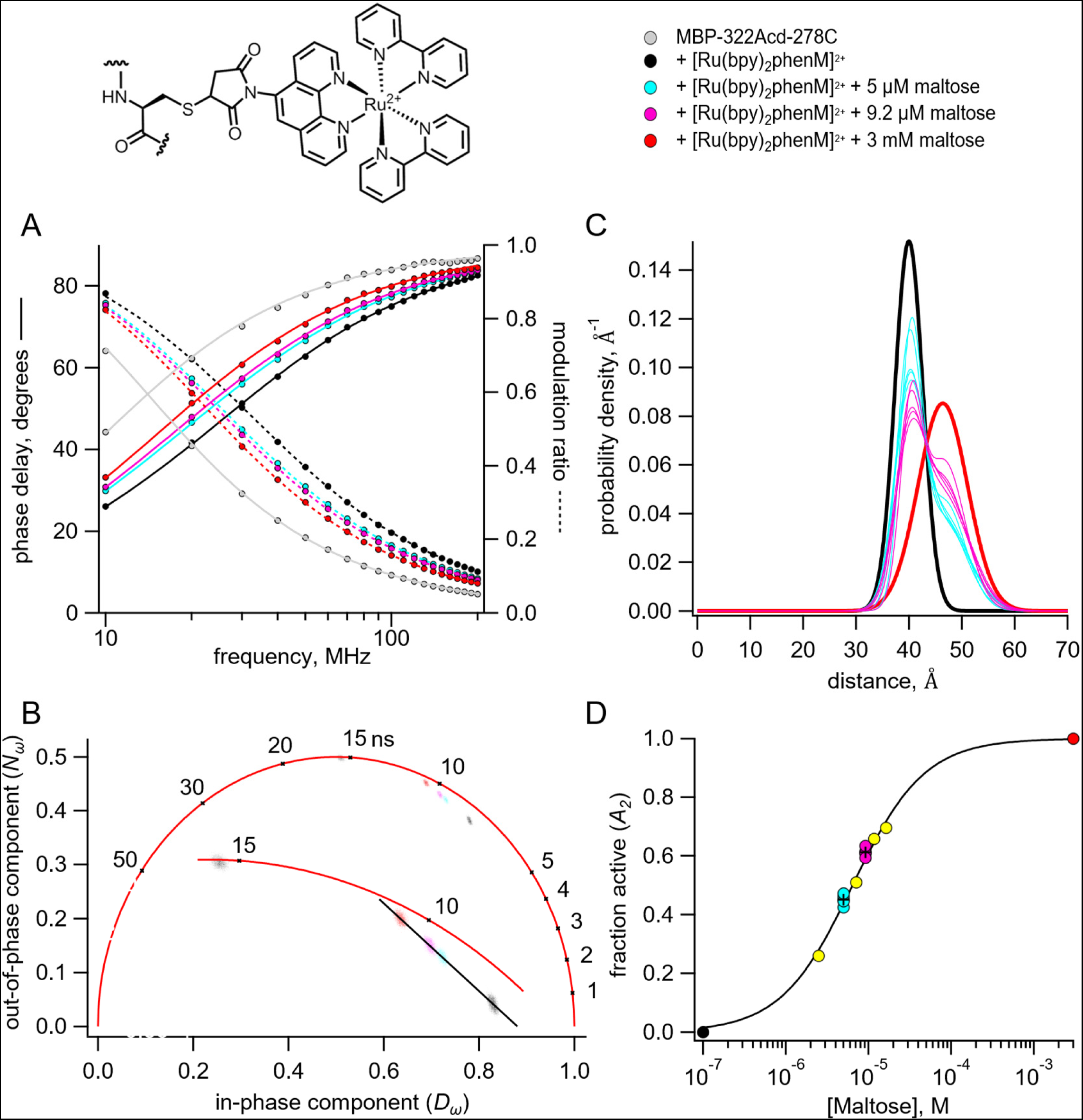
Time-resolved tmFRET of MBP-322Acd-278C labelled with [Ru(bpy)_2_phenM]^2+^ across a range of maltose concentrations. Structure of [Ru(bpy)_2_phenM]^2+^ and legend for all panels are shown at the top. (**A**) Weber plot of the fluorescence lifetimes with 0, 5 μM, 9.2 μM, and 3 mM maltose, in the presence and absence [Ru(bpy)_2_phenM]^2+^ (**B**) Phasor plot of the fluorescence lifetimes with 0, 5 μM, 9.2 μM, and 3 mM maltose, in the presence and absence of [Ru(bpy)_2_phenM]^2+^. The markers and numbers on the universal circle indicate time constants (in nanoseconds) for single exponential decays. A blowup of the region containing spots is shown in the inset. (**C**) Spaghetti plot of the distance distributions measured by time-resolved tmFRET with 5 μM, 9.2 μM (thin solid curves) and 0 and 3 mM (thick solid curves) maltose (n=5). (**D**) Scatter plot of maltose dose-response relation (n=5 for 5 μM and 9.2 μM maltose). The average values are shown as black +. A separate experiment with 5 different maltose concentrations is shown in yellow. Fit is with a Langmuir isotherm with a K_D_ of 6.3 μM maltose.

We also performed similar time-resolved FRET experiments on MBP-322Acd-278C with [Fe(phen maleimide)_3_]^2+^ ([Fe(phenM)_3_]^2+^). Like Ru^2+^, Fe^2+^ forms complexes with three bipyridyls or phenanthrolines that are highly absorbent in the visible range and can be used as FRET acceptors with visible fluorophores (11). The *R*_0_ of Acd with [Fe(phenM)_3_]^2^ is 41.8 Å. The addition of [Fe(phenM)_3_]^2+^ caused a dramatic decrease in the average fluorescence lifetime of MBP-322Acd-278C as expected for FRET between Acd and [Fe(phenM)_3_]^2+^ (Figure 4A). Furthermore, the average lifetime systematically increased upon addition of increasing concentrations of maltose, consistent with the maltose-dependent increase in distance predicted for MBP-322Acd-278C. No change in lifetime was observed for MBP-322Acd without the cysteine mutation (data not shown). Global fits of the Gaussian model to the data on the Weber plot at five different maltose concentrations (0, 5 μM, 9.2 μM, 12.9 μM, and 3 mM) yielded mean distances and standard deviations similar to those for [Ru(bpy)_2_phenM]^2+^, even though the *R_0_* was somewhat smaller (Figure 4D). The phasor plot (Figure 4B) and average distributions (Figure 4C) display a gradual shift in the equilibrium between the resting and active state with increasing maltose concentrations. Finally, the maltose dependence of the fraction of the active state nicely conformed to a binding isotherm, yielding an affinity of 9.2 μM for maltose (Figure 4E). These experiments demonstrate that both [Ru(bpy)_2_phenM]^2+^ and [Fe(phenM)_3_]^2+^ make good acceptors when using time-resolved tmFRET to determine distance distributions among conformational states and the free energy difference between conformational states.

**Figure 4.**
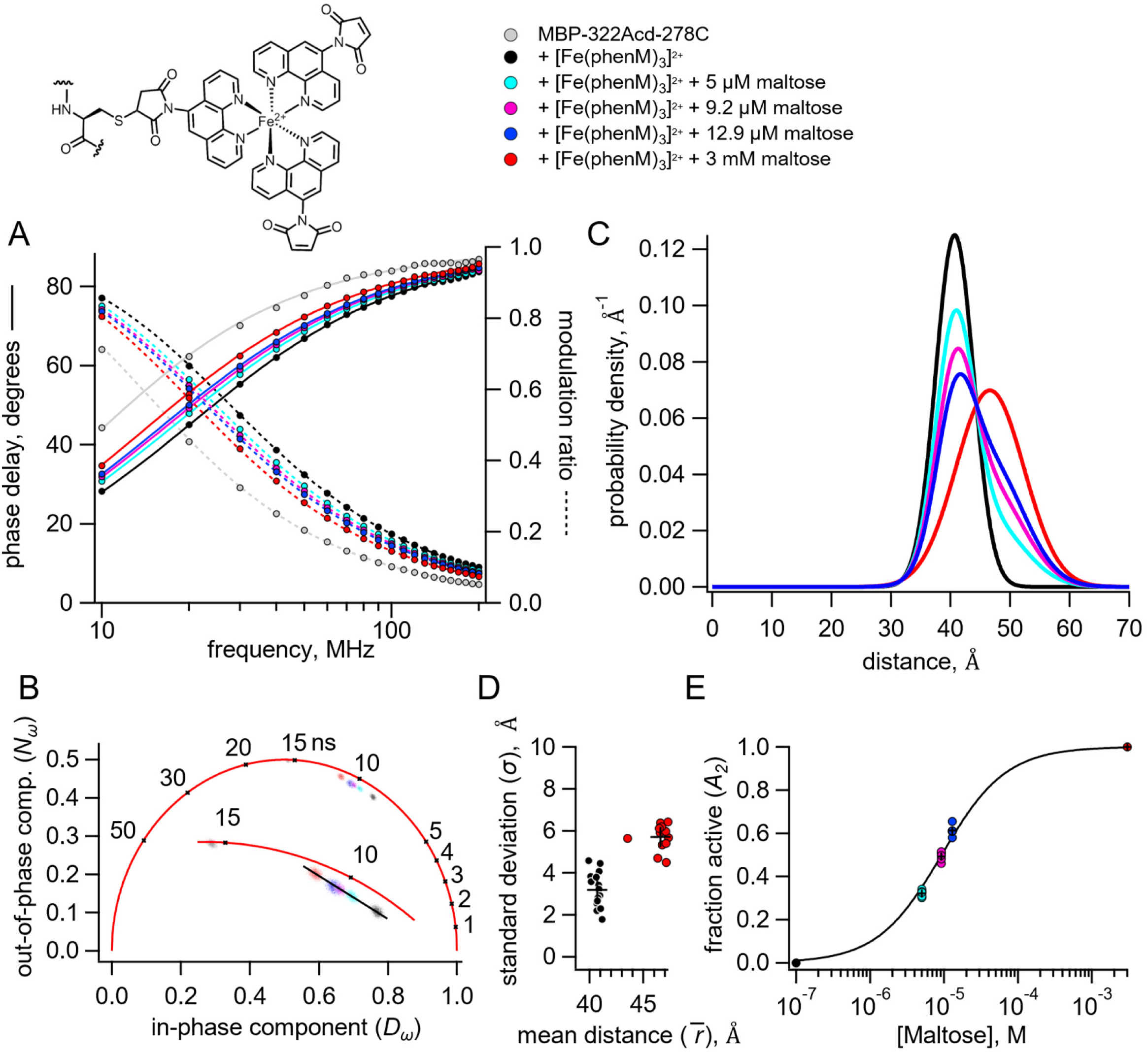
Time-resolved tmFRET of MBP-322Acd-278C labelled with [Fe(phenM)_3_]^2+^ across a range of maltose concentrations. Structure of [Fe(phenM)_3_]^2+^ and legend for all panels are shown at the top. (**A**) Weber plot of the fluorescence lifetimes with 0, 5 μM, 9.2 μM, 12.9 μM, and 3 mM, in the presence and absence of [Fe(phenM)_3_]^2+^. (**B**) Phasor plot of the fluorescence lifetimes with 0, 5 μM, 9.2 μM, 12.9 μM, and 3 mM, in the presence and absence of [Fe(phenM)_3_]^2+^. The markers and numbers on the universal circle indicate time constants (in nanoseconds) for single exponential decays. A blowup of the region containing spots is shown in the inset. (**C**) Plot of the average distance distributions measured by time-resolved tmFRET with 0, 5 μM, 9.2 μM, 12.9 μM, and 3 mM maltose (n=5). (**D**) Scatter plot of the standard deviation vs. the mean distance for the Gaussian distributions with 0 (n=16) and 3 mM (n=13) maltose. The average values are shown as black +. (**E**) Scatter plot of maltose dose-response relation (n=5 for 5 μM, 9.2 μM, 12.9 μM). The average values are shown as black +. Solid curve is a fit with a Langmuir isotherm with a K_D_ of 9.2 μM maltose.

**Figure 5.**
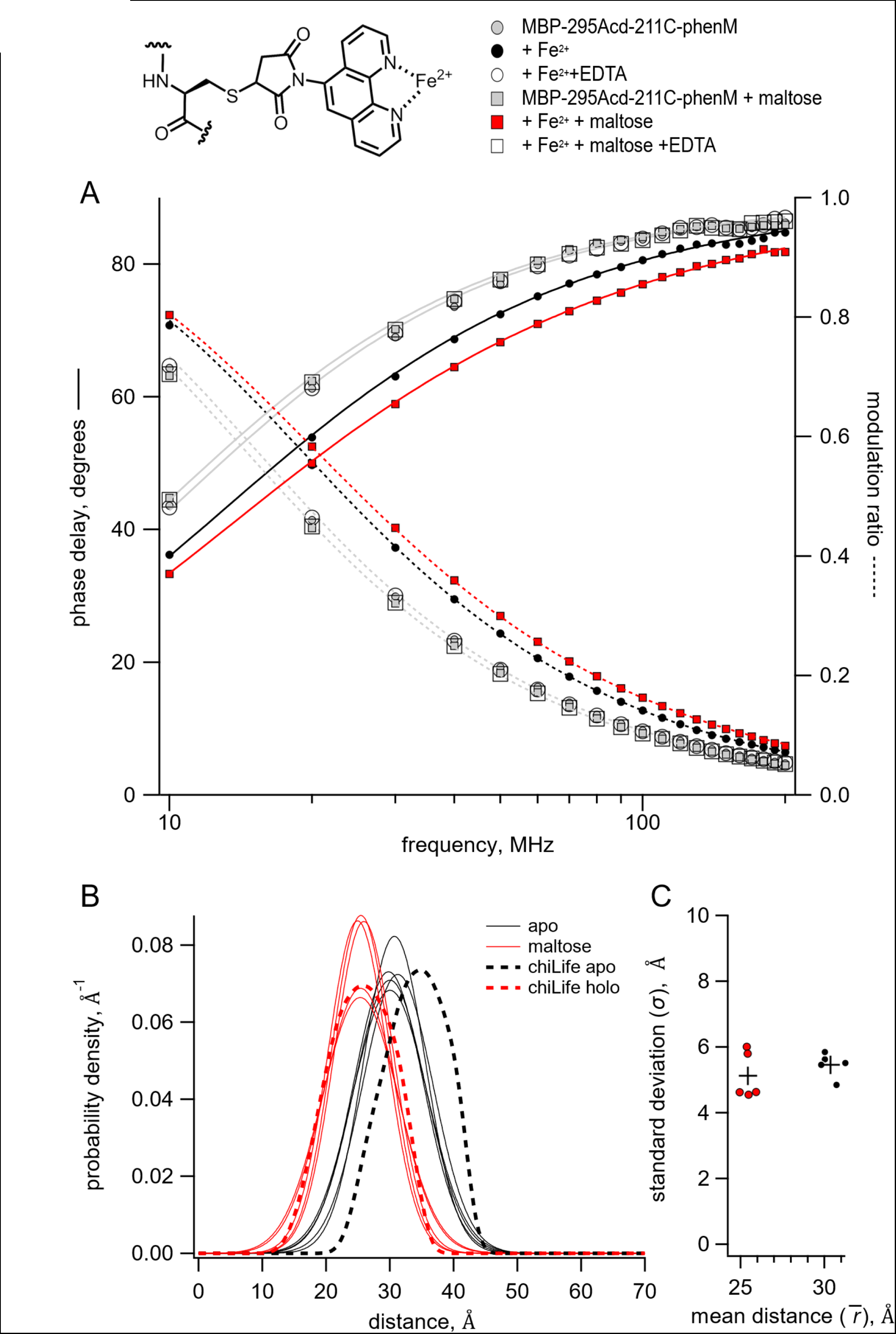
Time-resolved tmFRET of MBP-295Acd-211C labelled with [Fe(phenM)]^2+^. Structure of [Fe(phenM)]^2+^ and legend for all panels are shown at the top. (**A**) Weber plot of the fluorescence lifetimes of MBP-295Acd-211C-phenM with 0 and 3 mM maltose and in the presence and absence of Fe^2+^ and EDTA. (**B**) Spaghetti plot of the distance distributions measured by time-resolved tmFRET (thin solid curves) with 0 and 3 mM maltose (n=5) compared to the predicted distributions from chiLife for the resting and active state (dashed curves). (**C**) Scatter plot of the standard deviation vs. the mean distance for the Gaussian distributions with 0 and 3 mM maltose (n=5). The average values are shown as black +.

### Time-resolved FRET with [Fe(phenM)]^2+^

For MBP-295Acd-211C, the predicted donor-acceptor distances are shorter (on the order of 30 Å) and, instead of maltose increasing the donor-acceptor distance like for MBP-322Acd-278C, maltose decreases the distance. At these distances, the FRET efficiency with [Ru(bpy)_2_phenM]^2+^ and [Fe(phenM)_3_]^2+^ is nearly one, making these acceptors inadequate for detecting distance changes in MBP-295Acd-211C (11). However, we have found that Fe^2+^ bound to a single phenanthroline maleimide ([Fe(phenM)]^2+^) exhibits a much lower absorption and therefore a much lower *R*_0_ (24.4 Å), making it ideal for the donor-acceptor distances in MBP-295Acd-211C (11).

The addition of Fe^2+^ caused a dramatic decrease in the average fluorescence lifetime of MBP-295Acd-211C-phenM as expected for FRET between Acd and [Fe(phenM)]^2+^ (Figure 5A). The decrease in lifetime was even greater in the presence of saturating concentrations of maltose, as expected from the maltose-dependent decrease in distance between the donor and acceptor in MBP-295Acd-211C. For [Fe(phenM)]^2+^, but not for [Ru(bpy)_2_phenM]^2+^ or [Fe(phenM)_3_]^2+^, the FRET was fully reversible with EDTA, as seen by the open symbols in the Weber plot surrounding the original donor-only points (Figure 5A). Fitting the Gaussian model to the lifetime data revealed that MBP-295Acd-211C-phenM had a somewhat broader distance distribution than MBP-322Acd-278C, but with a maltose-dependent decrease in average distance (Figure 5B and C). Once again, the distance distributions were similar to those predicted by chiLife, albeit with a somewhat shorter distance than predicted for the apo state. These data indicate that [Fe(phenM)]^2+^ can be used as an effective FRET acceptor for lifetime measurements of distance distributions with mid-range donor-acceptor distances.

### Pulse dipolar EPR with [Cu(phenM)]^2+^

To further explore the heterogeneity contributed by the phenM side chain, we turned to pulse dipolar EPR spectroscopy. Pulse dipolar EPR methods such as double electron-electron resonance (DEER) and relaxation-induced dipolar modulation enhancement (RIDME) measure distance distributions between unpaired electrons introduced into proteins via site-directed spin labeling (36). These data are typically analyzed using non-parametric models and therefore place no underlying assumptions on the shape of the distance distributions, only that they are smooth. For these experiments, we introduced cysteine substitutions at both donor and acceptor sites in our previous MBP constructs, generating MBP-295C-211C and MBP-322C-278C. Nitroxide spin labels introduced at these site pairs have previously been employed to detect maltose-dependent conformational changes in MBP using DEER (13). To examine distance distributions obtained using the phenM label, MBP-295C-211C and MBP-322C-278C were labelled with phenM and subsequently labelled with Cu^2+^ (Figure 6A). Cu^2+^ is a d^9^ transition metal ion containing one unpaired electron (*S* = 1/2) and can therefore be used as a spin label for EPR experiments. Indeed, both MBP-295C-211C and MBP-322C-278C dual labeled with [Cu(phenM)]^2+^ displayed continuous-wave EPR spectra consistent with each Cu^2+^ coordinated by a single phenanthroline, indicating specific labeling of the phenM side chains with Cu^2+^ (Figure S4).

**Figure 6.**
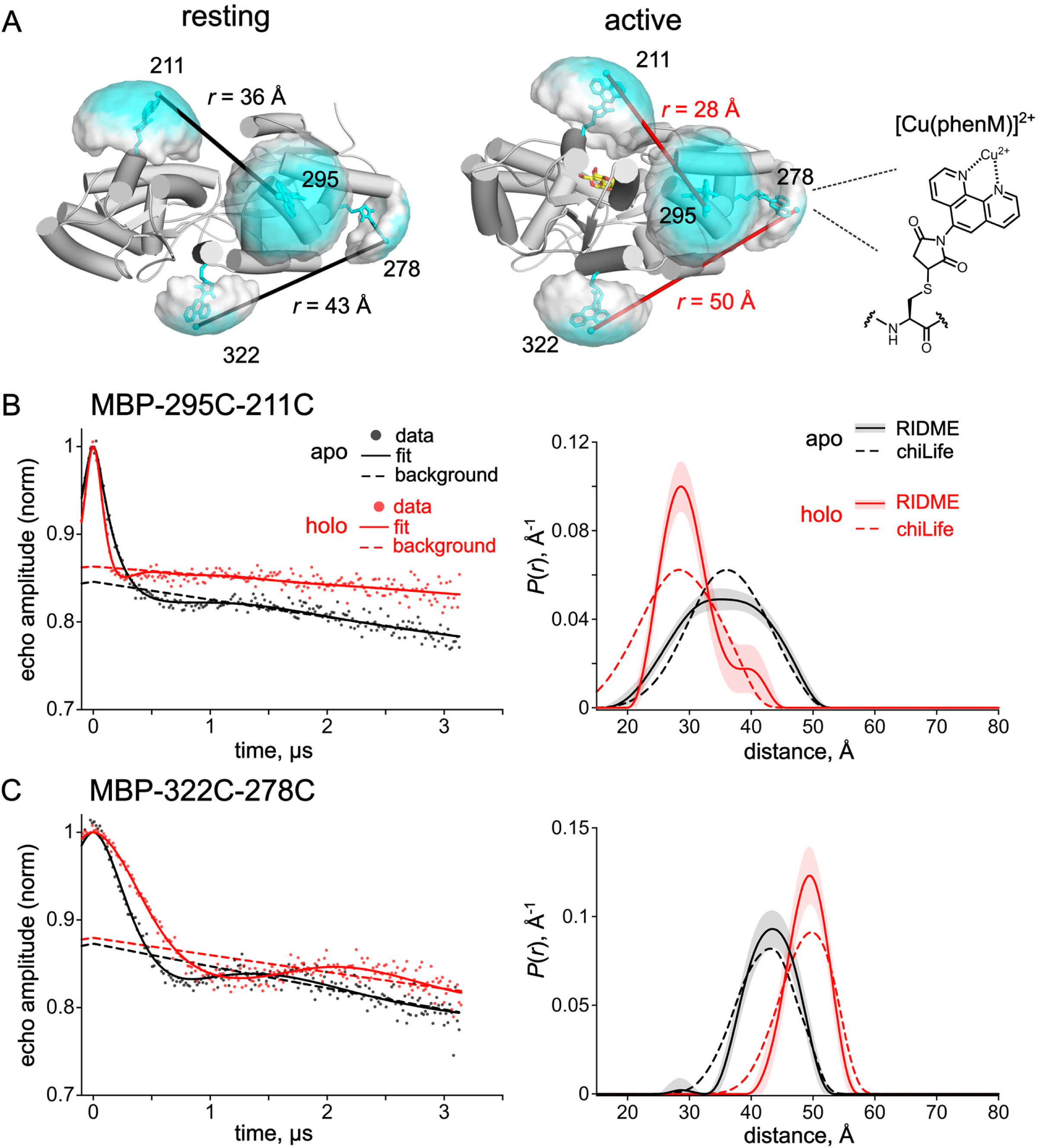
Pulse dipolar EPR distance distributions with the [Cu(phenM)]^2+^ spin label. (**A**) Cartoon representations of resting state (pdb 1omp) (35) and active state (pdb 1anf) (34) of MBP labeled *in silico* with [Cu(phenM)]^2+^ at 295C, 322C, 211C, and 278C using chiLife. Rotameric clouds are represented as surfaces and colored on a gradient from low (white) to high (cyan) probability. Predicted most probable distances determined by chiLife for the MBP-295C-211C and MBP-322C-278C site pairs are indicated. The structure of the spin-labeled [Cu(phenM)]^2+^ side chain is shown at right. (**B, C**) 5-pulse RIDME time-traces and calculated distance distributions for [Cu(phenM)]^2+^-labeled MBP-295C-211C (B) and MBP-322C-278C (C) in the absence of maltose (apo, black) and in the presence of 5 mM maltose (holo, red). Dashed curves on the distance distribution plot are predicted distributions calculated with chiLife. Shaded error bands on the RIDME distance distributions represent 95% confidence intervals determined from the DeerLab fits.

To determine distance distributions between [Cu(phenM)]^2+^ labels on MBP directly, we performed RIDME experiments in the absence and presence of saturating maltose (Figure 6B, S5). For MBP-295C-211C labeled with [Cu(phenM)]^2+^, RIDME in the absence of maltose reveals a broad distance distribution centered at 35.1 Å, which narrowed and shifted to a most probable distance of 28.6 Å in saturating maltose (Figure 6B). These distances are in excellent agreement with those predicted by chiLife (36.1 Å and 28.4 Å for apo and holo, respectively) (Figure 6B dashed line). RIDME data on MBP-322C-278C labeled with [Cu(phenM)]^2+^, positioned on the backside of the clamshell, reveal the expected increase in Cu^2+^‒Cu^2+^ distance upon addition of maltose, with distance distributions centered at 43.5 Å and 49.5 Å for apo and holo conditions, respectively (Figure 6C). Again, these distances are within 1 Å of the chiLife predictions (43.1 Å and 49.8 Å for apo and holo, respectively). These results establish [Cu-phenM]^2+^ as a useful and commercially available spin label for determining conformational distributions in proteins using pulse dipolar EPR spectroscopy. Moreover, the similarity of both the RIDME distributions (Figure 6B,C) and the tmFRET distributions (Figure 2B and 5B) to their respective chiLife predictions, and to each other, suggests the distributions at room temperature are well captured by the rapid freezing of the sample in the RIDME experiments. Overall, these experiments support the accuracy of RIDME, tmFRET, and chiLife for estimating the rotameric ensembles and resulting distance distributions involving [metal(phenM)]^2+^ labels.

## DISCUSSION

This paper applies time-resolved tmFRET to study protein allostery and conformational dynamics. tmFRET utilizes a fluorescent noncanonical amino acid as the donor and our new metal-bipyridyl derivatives as the acceptor to overcome limitations of traditional FRET methods (11). We applied this method to MBP and demonstrated it can accurately determine distances, conformational heterogeneity, and energetics. The results highlight the utility of time-resolved tmFRET in characterizing protein dynamics and conformational changes, offering valuable insights into the mechanisms of allosteric regulation.

In addition to its utility in tmFRET, we showed that the cysteine-reactive bipyridyl derivative phenM can be used with Cu^2+^ as a spin label in pulse dipolar EPR spectroscopy. Distance distributions from pulsed dipolar EPR are commonly determined using non-parametric models which have the advantage of requiring no underlying assumptions about the shape of the distance distribution. Using the same sites on MBP to which we introduced the donor and acceptor for tmFRET, we show that the distance distributions produced by RIDME were similar to the predictions of chiLife and also to the distance distributions measured with time-resolved tmFRET. These results both establish [Cu(phenM)]^2+^ as a spin label for pulse dipolar EPR spectroscopy and validate the distance distributions measured with time-resolved tmFRET.

The determination of distance distributions directly from the lifetime data is an ill-posed problem and, therefore, we needed to parameterize the distance distributions for our time-resolved tmFRET experiments. For our model, the heterogeneous distances between the donor and acceptor for each state were described by a Gaussian distribution with a distinct mean distance and standard deviation, although other distance distributions such as Lorentzians have also been used (37–39). Given these assumptions, values for the parameters were well determined from an analysis of the *χ*^2^ surfaces (Figure S2A-D) and were fairly consistent across multiple experiments (Figures 2C, 4D, 5C). In addition, the distances, conformational heterogeneity, and energetics determined from the Gaussian model were consistent with molecular modeling (chiLife, Figure 2B, 5B) and EPR spectroscopy (RIDME, Figure 6), and subsaturating maltose concentrations (Figure 3 and 4).

Despite our promising results, correlation of some parameters was observed and overfitting the data was a concern. For example, the value of standard deviations of the Gaussian distance distributions covaried with the fraction of donor only (Figure S2E,F), making it important that the amount of donor only is small (<20%) and is verified with independent experiments (see Figure S6 of (11)). Fitting a single condition (e.g., apo and holo) assuming two Gaussian components (two conformational states) did not always yield consistent results across experiments. For that reason, we assumed the apo and holo conditions contained only a single Gaussian component (a single conformation) for each condition. This is a reasonable assumption for MBP where the protein is thought to be largely in a single conformational state in apo and holo conditions (40). However, for many other proteins, we expect that apo and holo conditions will contain a mixture of resting and active states (Figure 1A) (41). Some future modifications that might help improve the fitting under these conditions include: 1) global fitting of the fluorescence lifetime and steady-state FRET efficiency under the same condition, 2) global fitting of the data across multiple similar FRET pairs with different *R*_0_ values, and 3) constraining one or more of the Gaussian components to the predictions of molecular modeling or molecular dynamics calculations. Ultimately, a Bayesian inference analysis of the Gaussian model is needed to characterize the uncertainties and correlations among the parameters, as has been done with DEER (42).

## AUTHOR CONTRIBUTIONS

WNZ, EGBE, and SEG designed experiments, performed research, analyzed data, and wrote the paper, PE performed research and analyzed data, MHT, EJP, and ST analyzed data, contributed tools, and edited the manuscript, and KDS contributed tools.

## ACKNOWLEDGEMENTS

We thank the Oregon State University GCE4ALL (Center for Genetic Code Expansion for All) for their longstanding collaboration, Dr. Shauna C. Otto (University of Washington) and Dr. Chloe Jones (University of Pennsylvania) for excellent technical support, and Richard W. Aldrich for everything else. Research reported in this publication was supported by the National Institutes of Health under award numbers R35GM145225 (to S.E.G.), R35GM148137 and R03TR004135 (to W.N.Z.), R01GM125753 (S.S.), T32EY007031 (to E.G.B,E), and T32GM008268 (to P.E). This research was also supported in part by the National Science Foundation under Grant DGE-1747486 (E.J.P.).

---

**Figure S1.**
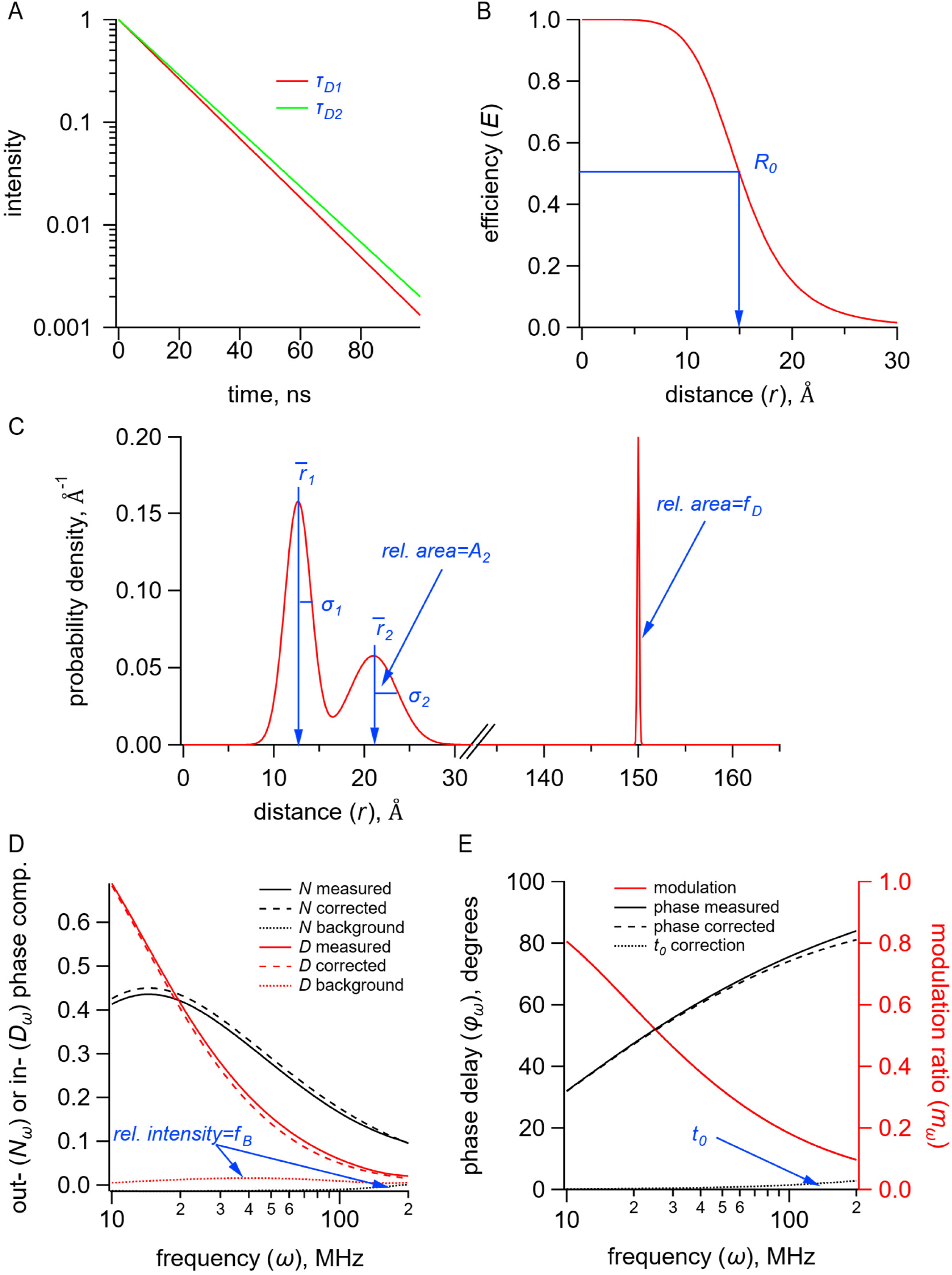
Gaussian model for time-resolved tmFRET. Parameters are in blue. (**A**) Plot of fluorescence lifetimes in the time domain for a single exponential donor fluorophore with time constants τ_D1_ and τ_D2_ in the resting and active states respectively. (**B**) Plot of the FRET efficiency (*E*) as a function of distance (*r*) showing the characteristic distance for the donor-acceptor pair (*R*_0_). (**C**) Plot of a distribution of donor-acceptor distances (*P*(*r*)) with two Gaussian components with means (*r̅*_1_ and *r̅*_2_), standard deviations (σ_1_ and σ_2_), and fraction active (*A*_2_). The fraction donor only (*f*_D_) was modeled as a narrow Gaussian with a mean distance of 150 Å and a standard deviation of 0.1 Å, too far to exhibit any detectable FRET. (**D**) Plot of the out-of-phase (*N*_*ω*_) and in-phase (*D*_*ω*_) components of the measured, corrected, and background fluorescence response as a function of the modulation frequency (ω) where *f*_B_is the fraction of the fluorescence intensity due to background. (**E**) Plot of the phase delay (*φ*_*ω*_) and modulation ratio (*m*_*ω*_) of the measured and corrected fluorescence response as a function of the modulation frequency (ω) where *t*_0_. is the time shift of the IRF.

**Figure S2.**
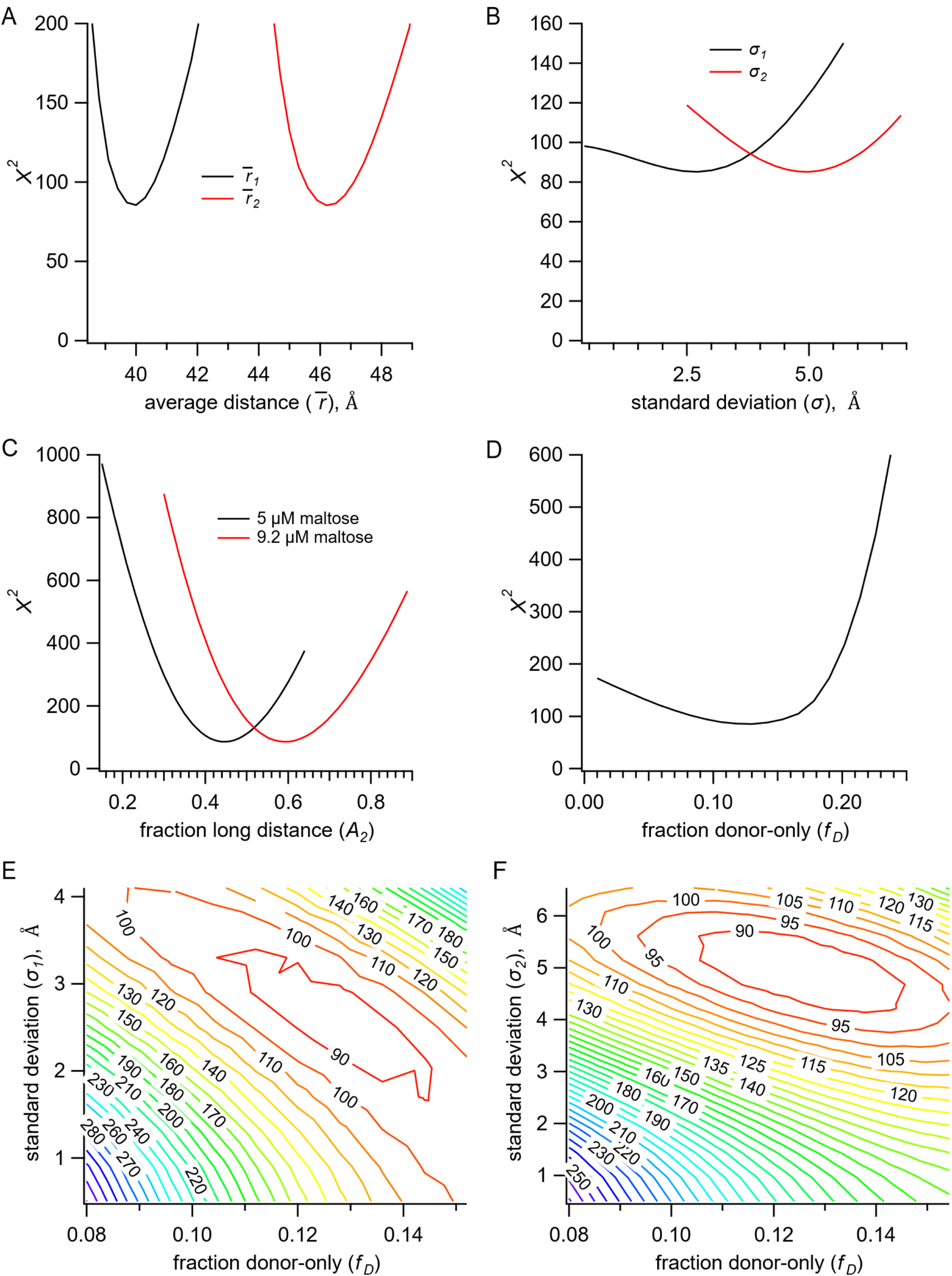
Identifiability of parameters in Gaussian model. (**A-D**) Plots of *χ*^2^ vs Gaussian means (*r̅*_1_ and *r̅*_2_) (A) and standard deviations (σ_1_ and σ_2_) (B), fraction active (*A*_2_) in 5 μM and 9.2 μM maltose (C), and fraction donor only (*f*_D_) (D). For each plot, minimum *χ*^2^was determined by global fitting with the single parameter at different fixed values, while all remaining parameters varied. Results are for a representative dataset from MBP-322Acd-278C with [Ru(bpy)_2_phenM]^2+^. (**E,F**) *χ*^2^surfaces for standard deviations (σ_1_ (E) and σ_2_ (F)) vs. fraction donor only (*f*_D_) showing some correlation between these parameters in the model. For each plot the standard deviation and fraction donor only were fixed at a range of values and the minimum *χ*^2^was determined by global fitting. Contour lines are labeled with the minimized *χ*^2^.

**Figure S3.**
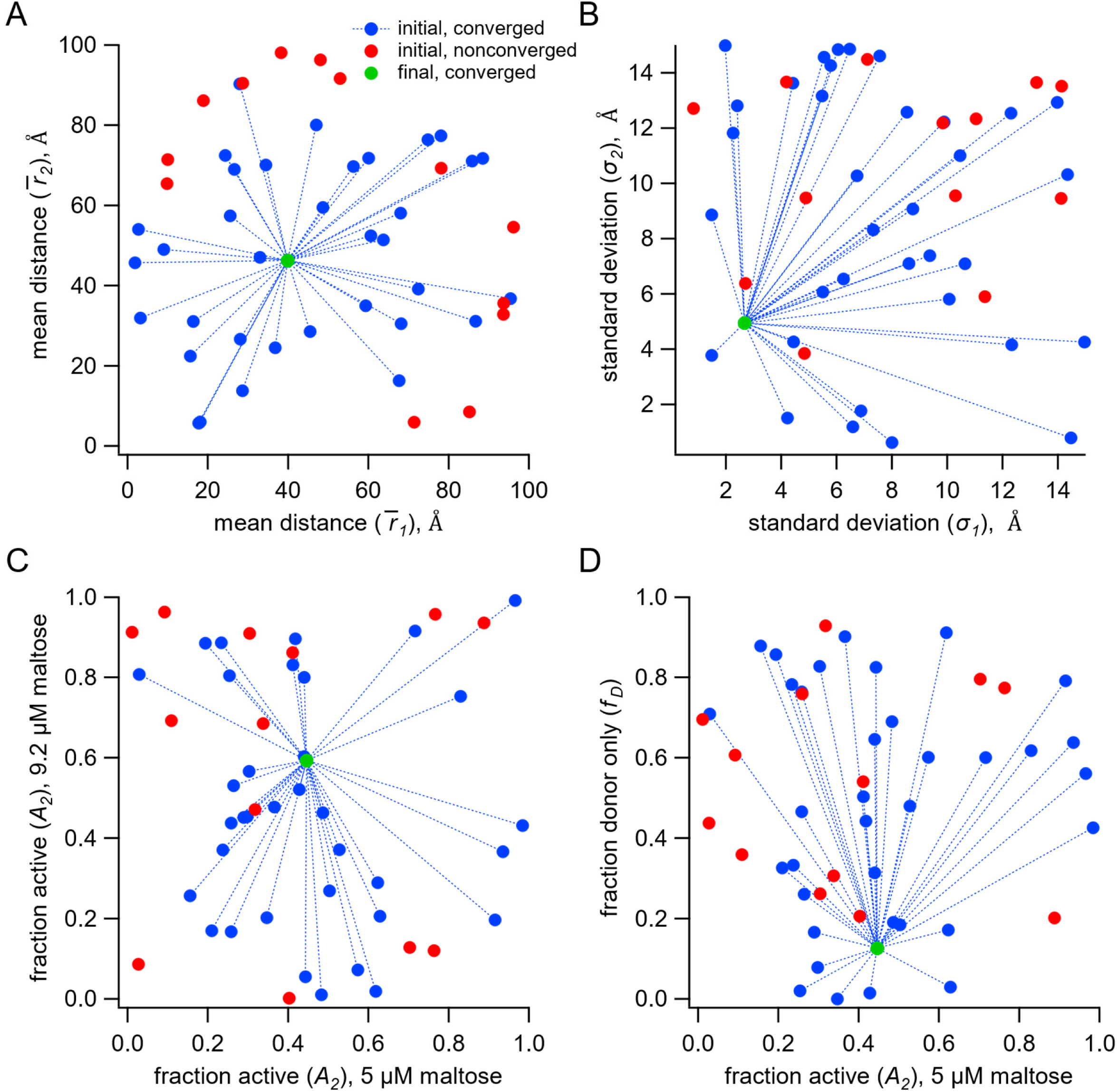
Convergence of the Gaussian model with different initial parameter values. (**A-D**) Plots of the initial values of parameters; *r̅*_1_ and *r̅*_2_ (A), σ_1_ and σ_2_ (B), *A*_2_ in 5 μM and 9.2 μM maltose (C), and *A*_2_ in 5 μM and *f*_D_(D); before convergence (blue circles) and after convergence (green circles) for different pairs of parameters. Initial conditions that did not converge or attain a minimum *χ*^2^less than 3000 are shown in red. For these calculations, the initial values for the shown parameters were varied randomly between the limits indicated by the axes, and global fitting was performed allowing all of the parameters to vary. All initial values in blue converged to nearly identical values for the parameters. Results are for a representative dataset from MBP-322Acd-278C with [Ru(bpy)_2_phenM]^2+^.

**Figure S4.**
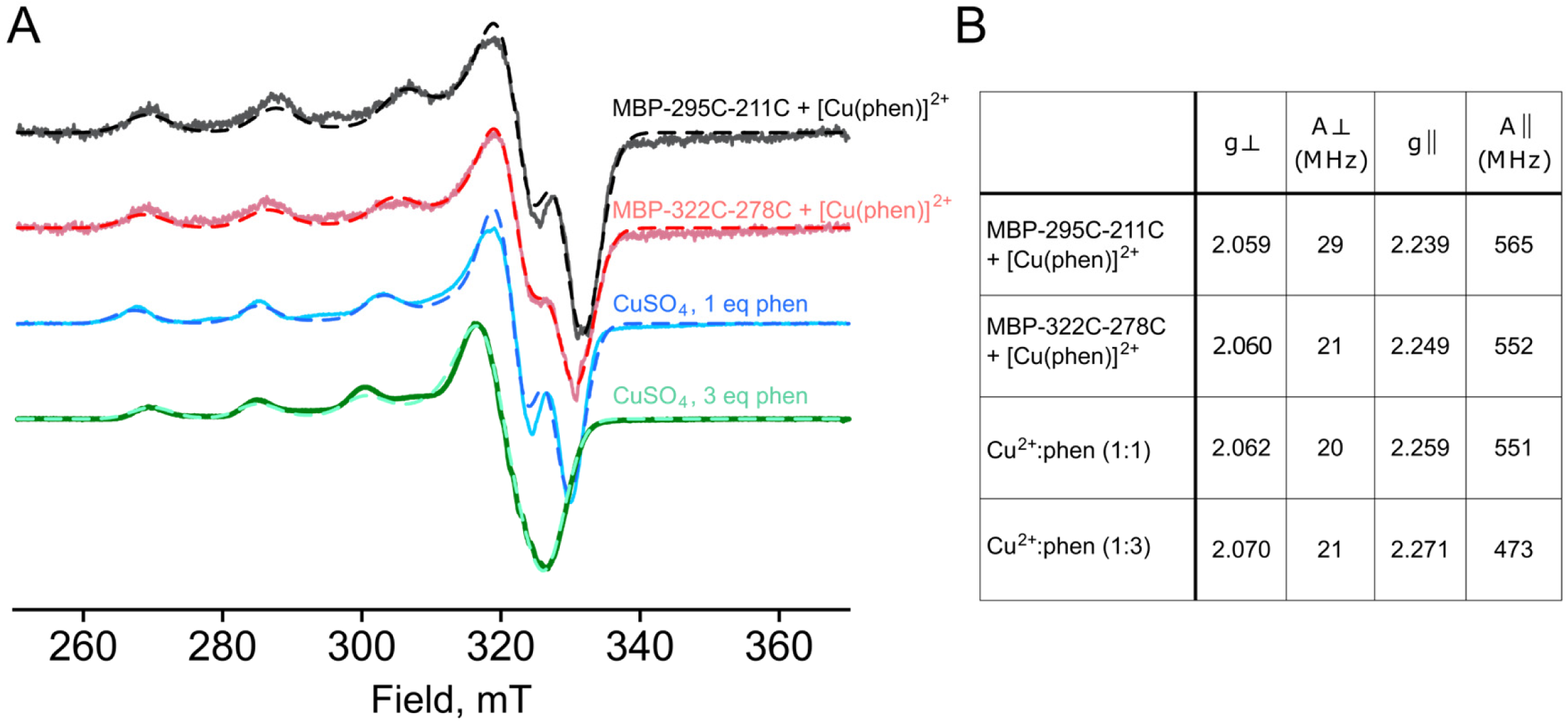
Cu^2+^ CW EPR. (**A**) X-band CW EPR spectra recorded at 112 K. Spectra are normalized by spectral intensity. Best fit simulations for each spectrum, performed in EasySpin, are shown as dashed curves. (**B**) Table of fitted magnetic parameters g and A from simulations shown in panel A.

**Figure S5.**
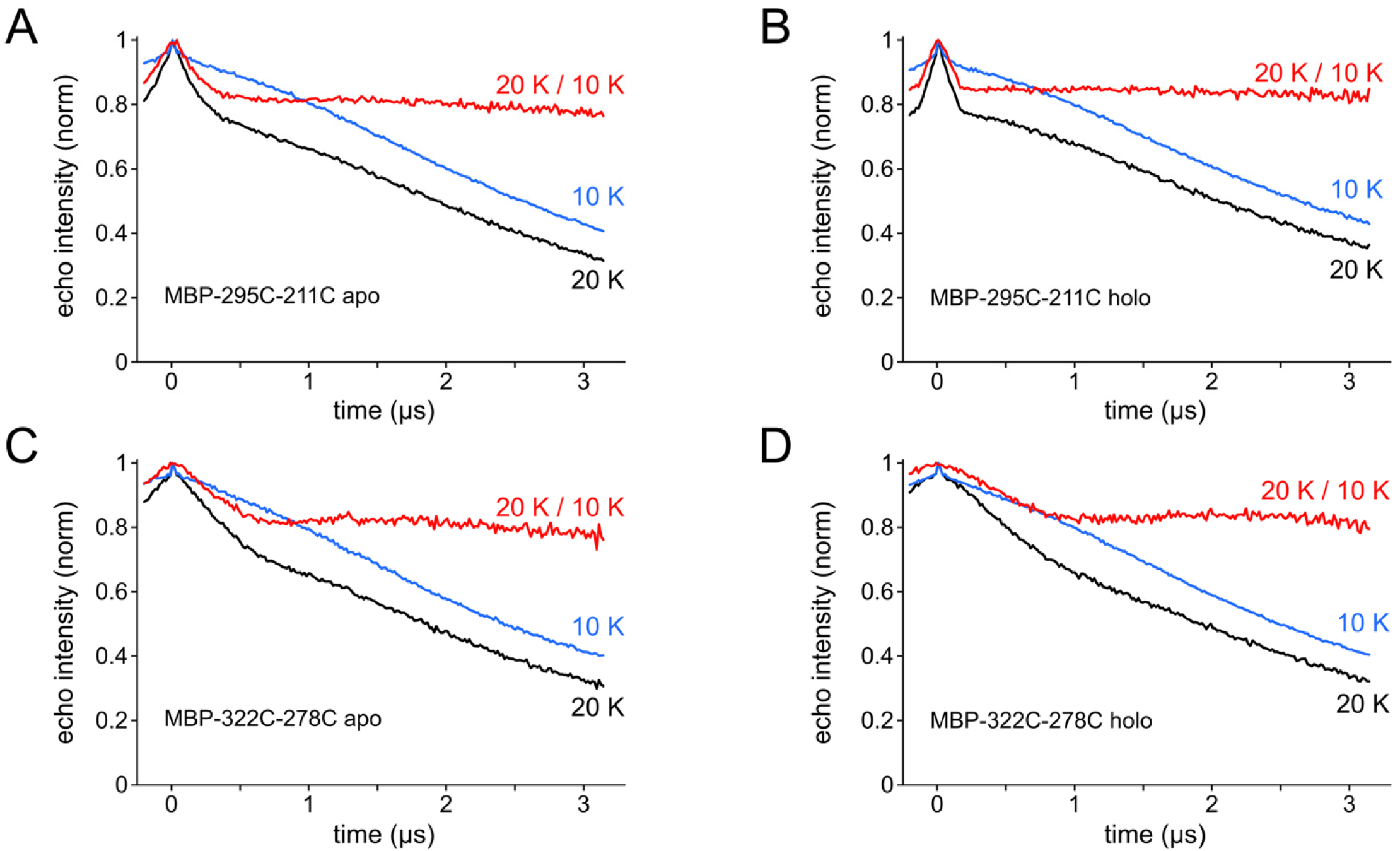
RIDME zero-time artifact removal by division. 5-pulse RIDME time-traces measured at 20 K (black) contained a sharp echo-crossing artifact at *t* ≈ 0 that was not removable through phase cycling. RIDME time-traces measured at 10 K (blue) also contain the zero-time artifact, but lack significant dipolar modulation, as the relaxation interval, *T*_R_, is only ∼ 4% of the Cu^2+^ *T*_1e_ at this temperature. Division of the 20 K data set by the 10 K data set removes the zero-time artifact, as well as much of the intermolecular background decay, but preserves the RIDME dipolar modulation (red trace). Data are shown for [Cu(phenM)]^2+^ labeled MBP-295C-211C apo (A), MBP-295C-211C holo (B), MBP-322C-278C apo (C), and MBP-322C-278C holo (D).

## REFERENCES

1. Stryer, L., and R. P. Haugland. 1967. Energy transfer: a spectroscopic ruler. Proc Natl Acad Sci U S A. 58(2):719–726, http://www.ncbi.nlm.nih.gov/pubmed/5233469.

2. Lakowicz, J. R. 2006. Principles of fluorescence spectroscopy. Springer, New York.

3. Eis, P. S., and D. P. Millar. 1993. Conformational distributions of a four-way DNA junction revealed by time-resolved fluorescence resonance energy transfer. Biochemistry. 32(50):13852–13860, doi: 10.1021/bi00213a014, https://www.ncbi.nlm.nih.gov/pubmed/8268160.

4. Hochstrasser, R. A., S. M. Chen, and D. P. Millar. 1992. Distance distribution in a dye-linked oligonucleotide determined by time-resolved fluorescence energy transfer. Biophys Chem. 45(2):133–141, doi: 10.1016/0301-4622(92)87005-4, https://www.ncbi.nlm.nih.gov/pubmed/1286148.

5. Yang, M., and D. P. Millar. 1996. Conformational flexibility of three-way DNA junctions containing unpaired nucleotides. Biochemistry. 35(24):7959–7967, doi: 10.1021/bi952892z, https://www.ncbi.nlm.nih.gov/pubmed/8672499.

6. Gryczynski, I., W. Wiczk, M. L. Johnson, H. C. Cheung, C. K. Wang, and J. R. Lakowicz. 1988. Resolution of end-to-end distance distributions of flexible molecules using quenching-induced variations of the Forster distance for fluorescence energy transfer. Biophys J. 54(4):577–586, doi: 10.1016/S0006-3495(88)82992-8, https://www.ncbi.nlm.nih.gov/pubmed/3224143.

7. Lakowicz, J. R., I. Gryczynski, H. C. Cheung, C. K. Wang, and M. L. Johnson. 1988. Distance distributions in native and random-coil troponin I from frequency-domain measurements of fluorescence energy transfer. Biopolymers. 27(5):821–830, doi: 10.1002/bip.360270509, https://www.ncbi.nlm.nih.gov/pubmed/3382720.

8. Lakowicz, J. R., I. Gryczynski, W. Wiczk, G. Laczko, F. C. Prendergast, and M. L. Johnson. 1990. Conformational distributions of melittin in water/methanol mixtures from frequency-domain measurements of nonradiative energy transfer. Biophys Chem. 36(2):99–115, doi: 10.1016/0301-4622(90)85014-w, https://www.ncbi.nlm.nih.gov/pubmed/2207280.

9. Lakowicz, J. R., J. Kuśba, W. Wiczk, I. Gryczynski, H. Szmacinski, and M. L. Johnson. 1991. Resolution of the conformational distribution and dynamics of a flexible molecule using frequency-domain fluorometry. Biophys Chem. 39(1):79–84, doi: 10.1016/0301-4622(91)85008-e, https://www.ncbi.nlm.nih.gov/pubmed/2012836.

10. Zagotta, W. N., B. S. Sim, A. K. Nhim, M. M. Raza, E. G. Evans, Y. Venkatesh, C. M. Jones, R. A. Mehl, E. J. Petersson, and S. E. Gordon. 2021. An improved fluorescent noncanonical amino acid for measuring conformational distributions using time-resolved transition metal ion FRET. Elife. 10, doi: 10.7554/eLife.70236, https://www.ncbi.nlm.nih.gov/pubmed/34623258.

11. Gordon, S. E., E. G. B. Evans, S. C. Otto, M. H. Tessmer, K. D. Shaffer, M. T. Gordon, E. J. Petersson, S. Stoll, and W. N. Zagotta. 2023. Long-distance tmFRET using bipyridyl-and phenanthroline-based ligands. Submitted.

12. Sungwienwong, I., Z. M. Hostetler, R. J. Blizzard, J. J. Porter, C. M. Driggers, L. Z. Mbengi, J. A. Villegas, L. C. Speight, J. G. Saven, J. J. Perona, R. M. Kohli, R. A. Mehl, and E. J. Petersson. 2017. Improving target amino acid selectivity in a permissive aminoacyl tRNA synthetase through counter-selection. Org Biomol Chem. 15(17):3603–3610, doi: 10.1039/c7ob00582b, https://www.ncbi.nlm.nih.gov/pubmed/28397914.

13. Jana, S., E. G. B. Evans, H. S. Jang, S. Zhang, H. Zhang, A. Rajca, S. E. Gordon, W. N. Zagotta, S. Stoll, and R. A. Mehl. 2023. Ultrafast Bioorthogonal Spin-Labeling and Distance Measurements in Mammalian Cells Using Small, Genetically Encoded Tetrazine Amino Acids. J Am Chem Soc. doi: 10.1021/jacs.3c00967, https://www.ncbi.nlm.nih.gov/pubmed/37364003.

14. Digman, M. A., V. R. Caiolfa, M. Zamai, and E. Gratton. 2008. The phasor approach to fluorescence lifetime imaging analysis. Biophys J. 94(2):L14–16, doi: 10.1529/biophysj.107.120154, https://www.ncbi.nlm.nih.gov/pubmed/17981902.

15. Colyer, R. A., C. Lee, and E. Gratton. 2008. A novel fluorescence lifetime imaging system that optimizes photon efficiency. Microsc Res Tech. 71(3):201–213, doi: 10.1002/jemt.20540, https://www.ncbi.nlm.nih.gov/pubmed/18008362.

16. Lakowicz, J. R., I. Gryczynski, G. Laczko, W. Wiczk, and M. L. Johnson. 1994. Distribution of distances between the tryptophan and the N-terminal residue of melittin in its complex with calmodulin, troponin C, and phospholipids. Protein Sci. 3(4):628–637, doi: 10.1002/pro.5560030411, https://www.ncbi.nlm.nih.gov/pubmed/8003981.

17. Cheung, H. C., I. Gryczynski, H. Malak, W. Wiczk, M. L. Johnson, and J. R. Lakowicz. 1991. Conformational flexibility of the Cys 697-Cys 707 segment of myosin subfragment-1. Distance distributions by frequency-domain fluorometry. Biophys Chem. 40(1):1–17, doi: 10.1016/0301-4622(91)85025-l, https://www.ncbi.nlm.nih.gov/pubmed/1873469.

18. Tessmer, M. H., and S. Stoll. 2023. chiLife: An open-source Python package for in silico spin labeling and integrative protein modeling. PLoS Comput Biol. 19(3):e1010834, doi: 10.1371/journal.pcbi.1010834, https://www.ncbi.nlm.nih.gov/pubmed/37000838.

19. Tessmer, M. H., E. R. Canarie, and S. Stoll. 2022. Comparative evaluation of spin-label modeling methods for protein structural studies. Biophys J. 121(18):3508–3519, doi: 10.1016/j.bpj.2022.08.002, https://www.ncbi.nlm.nih.gov/pubmed/35957530.

20. Hagelueken, G., R. Ward, J. H. Naismith, and O. Schiemann. 2012. MtsslWizard: In Silico Spin-Labeling and Generation of Distance Distributions in PyMOL. Appl Magn Reson. 42(3):377–391, doi: 10.1007/s00723-012-0314-0, https://www.ncbi.nlm.nih.gov/pubmed/22448103.

21. Spicher, S., and S. Grimme. 2020. Robust Atomistic Modeling of Materials, Organometallic, and Biochemical Systems. Angew Chem Int Ed Engl. 59(36):15665–15673, doi: 10.1002/anie.202004239, https://www.ncbi.nlm.nih.gov/pubmed/32343883.

22. Stoll, S., and A. Schweiger. 2006. EasySpin, a comprehensive software package for spectral simulation and analysis in EPR. J Magn Reson. 178(1):42–55, doi: 10.1016/j.jmr.2005.08.013, https://www.ncbi.nlm.nih.gov/pubmed/16188474.

23. Milikisyants, S., F. Scarpelli, M. G. Finiguerra, M. Ubbink, and M. Huber. 2009. A pulsed EPR method to determine distances between paramagnetic centers with strong spectral anisotropy and radicals: the dead-time free RIDME sequence. J Magn Reson. 201(1):48–56, doi: 10.1016/j.jmr.2009.08.008, https://www.ncbi.nlm.nih.gov/pubmed/19758831.

24. Abdullin, D., M. Suchatzki, and O. Schiemann. 2022. Six-Pulse RIDME Sequence to Avoid Background Artifacts. Appl. Magn. Reson. 53:539–554, doi: 10.1007/s00723-021-01326-1.

25. Ritsch, I., H. Hintz, G. Jeschke, A. Godt, and M. Yulikov. 2019. Improving the accuracy of Cu(ii)-nitroxide RIDME in the presence of orientation correlation in water-soluble Cu(ii)-nitroxide rulers. Phys Chem Chem Phys. 21(19):9810–9830, doi: 10.1039/c8cp06573j, https://www.ncbi.nlm.nih.gov/pubmed/31025988.

26. Fábregas Ibáñez, L., G. Jeschke, and S. Stoll. 2020. DeerLab: a comprehensive software package for analyzing dipolar electron paramagnetic resonance spectroscopy data. Magn Reson (Gott). 1(2):209–224, doi: 10.5194/mr-1-209-2020, https://www.ncbi.nlm.nih.gov/pubmed/34568875.

27. Fábregas-Ibáñez, L., G. Jeschke, and S. Stoll. 2022. Compactness regularization in the analysis of dipolar EPR spectroscopy data. J Magn Reson. 339:107218, doi: 10.1016/j.jmr.2022.107218, https://www.ncbi.nlm.nih.gov/pubmed/35439683.

28. Jones, C. M., Y. Venkatesh, and E. J. Petersson. 2020. Protein labeling for FRET with methoxycoumarin and acridonylalanine. Methods Enzymol. 639:37–69, doi: 10.1016/bs.mie.2020.04.008, https://www.ncbi.nlm.nih.gov/pubmed/32475410.

29. Gordon, S. E., E. G. B. Evans, S. C. Otto, E. J. Petersson, K. D. Shaffer, M. T. Gordon, M. H. Tessmer, S. Stoll, and W. N. Zagotta. 2023. Long-distance tmFRET using bipyridyl-and phenanthroline-based ligands. Submitted.

30. Speight, L. C., A. K. Muthusamy, J. M. Goldberg, J. B. Warner, R. F. Wissner, T. S. Willi, B. F. Woodman, R. A. Mehl, and E. J. Petersson. 2013. Efficient synthesis and in vivo incorporation of acridon-2-ylalanine, a fluorescent amino acid for lifetime and Förster resonance energy transfer/luminescence resonance energy transfer studies. J Am Chem Soc. 135(50):18806–18814, doi: 10.1021/ja403247j, https://www.ncbi.nlm.nih.gov/pubmed/24303933.

31. Haas, E., E. Katchalski-Katzir, and I. Z. Steinberg. 1978. Effect of the orientation of donor and acceptor on the probability of energy transfer involving electronic transitions of mixed polarization. Biochemistry. 17(23):5064–5070, doi: 10.1021/bi00616a032, https://www.ncbi.nlm.nih.gov/pubmed/718874.

32. Haas, E., M. Wilchek, E. Katchalski-Katzir, and I. Z. Steinberg. 1975. Distribution of end-to-end distances of oligopeptides in solution as estimated by energy transfer. Proc Natl Acad Sci U S A. 72(5):1807–1811, doi: 10.1073/pnas.72.5.1807, https://www.ncbi.nlm.nih.gov/pubmed/1057171.

33. Grinvald, A., E. Haas, and I. Z. Steinberg. 1972. Evaluation of the distribution of distances between energy donors and acceptors by fluorescence decay. Proc Natl Acad Sci U S A. 69(8):2273–2277, doi: 10.1073/pnas.69.8.2273, https://www.ncbi.nlm.nih.gov/pubmed/16592008.

34. Quiocho, F. A., J. C. Spurlino, and L. E. Rodseth. 1997. Extensive features of tight oligosaccharide binding revealed in high-resolution structures of the maltodextrin transport/chemosensory receptor. Structure. 5(8):997–1015, doi: 10.1016/s0969-2126(97)00253-0, https://www.ncbi.nlm.nih.gov/pubmed/9309217.

35. Sharff, A. J., L. E. Rodseth, J. C. Spurlino, and F. A. Quiocho. 1992. Crystallographic evidence of a large ligand-induced hinge-twist motion between the two domains of the maltodextrin binding protein involved in active transport and chemotaxis. Biochemistry. 31(44):10657–10663, doi: 10.1021/bi00159a003, https://www.ncbi.nlm.nih.gov/pubmed/1420181.

36. Goldfarb, D. 2022. Exploring protein conformations in vitro and in cell with EPR distance measurements. Curr Opin Struct Biol. 75:102398, doi: 10.1016/j.sbi.2022.102398, https://www.ncbi.nlm.nih.gov/pubmed/35667279.

37. Wu, P., and L. Brand. 1994. Conformational flexibility in a staphylococcal nuclease mutant K45C from time-resolved resonance energy transfer measurements. Biochemistry. 33(34):10457–10462, doi: 10.1021/bi00200a029, https://www.ncbi.nlm.nih.gov/pubmed/8068683.

38. Amir, D., and E. Haas. 1986. Determination of intramolecular distance distributions in a globular protein by nonradiative excitation energy transfer measurements. Biopolymers. 25(2):235–240, doi: 10.1002/bip.360250205, https://www.ncbi.nlm.nih.gov/pubmed/2420384.

39. Amir, D., S. Krausz, and E. Haas. 1992. Detection of local structures in reduced unfolded bovine pancreatic trypsin inhibitor. Proteins. 13(2):162–173, doi: 10.1002/prot.340130210, https://www.ncbi.nlm.nih.gov/pubmed/1377825.

40. Tang, C., C. D. Schwieters, and G. M. Clore. 2007. Open-to-closed transition in apo maltose-binding protein observed by paramagnetic NMR. Nature. 449(7165):1078–1082, doi: 10.1038/nature06232, https://www.ncbi.nlm.nih.gov/pubmed/17960247.

41. DeBerg, H. A., P. S. Brzovic, G. E. Flynn, W. N. Zagotta, and S. Stoll. 2016. Structure and Energetics of Allosteric Regulation of HCN2 Ion Channels by Cyclic Nucleotides. J Biol Chem. 291(1):371–381, doi: 10.1074/jbc.M115.696450, https://www.ncbi.nlm.nih.gov/pubmed/26559974.

42. Sweger, S. R., S. Pribitzer, and S. Stoll. 2020. Bayesian Probabilistic Analysis of DEER Spectroscopy Data Using Parametric Distance Distribution Models. J Phys Chem A. 124(30):6193–6202, doi: 10.1021/acs.jpca.0c05026, https://www.ncbi.nlm.nih.gov/pubmed/32614584.

